# A Root Mean Square Deviation Estimation Algorithm (REA) and its use for improved RNA Structure Prediction

**DOI:** 10.1101/2024.02.28.582508

**Authors:** Agoritsa Kalampaliki, Alexandros C. Dimopoulos, Martin Reczko

**Affiliations:** Institute for Fundamental Biomedical Science, Biomedical Sciences Research Center “Alexander Fleming”, 16672 Vari, Greece; Hellenic Naval Academy, 18539 Piraeus, Greece

## Abstract

The 3D structure of RNA is crucial for biotechnological applications and to comprehend its biological function. Recent developments using AlphaFold-inspired deep neural networks improved the prediction of 3D structure from RNA sequence, but evaluation of the accuracy of these predictions is still necessary. We present the RMSD Estimation Algorithm (REA), a feed-forward neural network to predict the root-mean-square deviation (RMSD) of a 3D RNA structure from its experimentally determined counterpart using its Molprobity [1] stereochemical validation features. It is trained on structures predicted by the DeepFoldRNA [2] and trRosettaRNA [3] transformer-based deep neural networks on a set of 182 models of RNA structures with pseudoknots. We compare REA with ARES [4], a deep learning algorithm that predicts the RMSD by extracting geometric patterns with equivariant convolution, assessing the prediction accuracy on RNAs with and without pseudoknots. REA outperformed ARES on both test sets with smaller absolute difference between the true and the predicted RMSD. Using a combination of REA and a Support Vector Regression (SVR) trained on the same data as REA, we can select RNA structures predicted with DeepFoldRNA, trRosettaRNA and Rhofold [5] to achieve a significantly higher prediction accuracy than any of the prediction methods used alone. This was shown on a validation set with 261 novel RNA chains extracted from the Nonredundant 3D Structure Dataset [5] and a test set with 55 novel RNA chains from RNA-Puzzles [5]. Our selection based prediction method can easily incorporate additional prediction algorithms.

## Introduction

RNA molecules, particularly non-coding RNAs, have gained significant attention from the scientific community due to their various roles in cells. The protein-coding RNA is the mRNA (messenger RNA) that also folds cotranscriptionally with many implications for gene expression regulation [6], while the rest of the RNA types, including tRNA (transfer RNA), rRNA (ribosomal RNA), miRNA (microRNA), siRNA (small interfering RNA), lncRNA (long non-coding RNA) belong to the non-coding RNAs category [7]. Non-coding RNAs have been discovered to have regulatory and enzymatic functions such as ribozymes, riboswitches, and transcription regulation [8]. Consequently, understanding the functions of RNA is crucial for studying cellular processes, but also for combating viral infections caused by retroviruses, such as HIV and SARS-CoV2 [8].

RNA molecules are chains consisting of the nucleotides adenine (A), uracil (U), guanine (G) and cytosine (C). Nucleotides within an RNA molecule can form chemical interactions with each other, leading to the molecule adopting a specific 3D structure. The most common type of interaction between nucleotides in RNA is the formation of Watson-Crick base pairs, which contribute to the formation of a helical structure [8]. The presence or absence of Watson-Crick base pairs between nucleotides defines the secondary structure of RNA. Apart from the Watson-Crick base pairs there are additional types of interactions between nucleotides in RNA such as non-Watson-Crick base pairs, base-backbone interactions, base stacking and long-range Watson-Crick base pairs (pseudoknots) that are included in an augmented secondary structure representation [8]. A pseudoknot is a nucleic acid secondary structure containing at least two stem-loop structures in which along the sequence half of one stem is inserted between the two halves of another stem. These various interactions collectively contribute to the overall 3D structure of RNA.

Determining the 2D structure of the RNA is the first step for understanding the RNA structure. Computational methods such as PETfold [9] and SPOT-RNA [10], a deep learning method, can predict an estimate for the 2D structure of RNA from sequence. Experimental methods have also been developed for determining presence of nucleotides in helices in the RNA molecule with the techniques of high-throughput sequencing and probing [11].

Although the secondary structure concerns mainly the helices, important parts of the 3D structure are loops and bulges where in most cases locations without Watson-Crick base pairs are found. Loops, located at the ends or between helices, can influence the orientation of helices and adopt specific geometric motifs that serve functional purposes [8]. Pseudoknots are also important in RNA 3D structure as they play an important role in enhancing the stability of the RNA structure [8]. The structure of loops, bulges and pseudoknots add to the secondary structure of RNA and they are necessary for the complete representation of RNA 3D structure.

Sequencing experiments have provided the sequences of multiple non-coding RNAs, which are stored in the Rfam database [12] and RNAcentral [13], along with their predicted secondary structures. RNA 3D structure is experimentally determined with x-ray crystallography, NMR or cryo-electron microscopy (cryo-EM). There are RNA structures from various organisms, complexed with proteins, nucleotide acids, and ligands are stored in the Protein Data Bank [14] at RCSB.org. Interestingly, cryo-electron microscopy (cryo-EM) data is being increasingly used to determine RNA structures, capturing conformational ensembles of RNAs, alternative structures of the same RNA captured in one sample [8].

The abundance of sequencing data and at the same time the difficulty of the experiments for structure determination push scientists towards predicting the 3D structure of RNA from sequence with computational methods. The ratio of RNA families (the RNAs with common functionality that have similar structure and sequence ex. rRNA, miRNA, lncRNA ect.) that are represented in the PDB is 125 / 4000, which shows that there are RNA types that don’t have an experimentally determined structure [8]. Additionally, the number of RNA structures in PDB is significantly smaller than the number of protein structures. As of 15 June 2023 there are 201885 structures containing a protein but only 6907 structures containing a RNA [15]. The small number of RNA structures available increase the difficulty of predicting the RNA 3D structure by limiting the number of structural homologs for homology modeling (starting from the sequence of an RNA and a homolog 3D structure to predict the 3D structure of the starting sequence) [8], limiting the number of training examples for machine learning approaches and by providing limited options of new and complex structures for blind assessment of modeling methods [16]. Until today, the sequence-based RNA 3D structure prediction, reviewed in [17], remains a challenge [16].

RNA-Puzzles are a series of community-wide experiments for the prediction of RNA 3D structures from sequence [18]–[21]. The latest RNA-Puzzles took place in collaboration with CASP15 - a regular benchmark event traditionally evaluating the protein structure prediction systems - and the assessment of the predicted models is published as a preprint [16]. The evaluation of the predicted structures focused on the topology of the predicted structure compared to the experimentally determined structure, but also considered the local quality and clashes in the predicted structure. The recent RNA-Puzzles challenge included 6 RNA-only targets, 4 non-natural RNA targets that are designed by humans and not found in nature, and 2 RNA-protein complexes targets. Of all targets the 2 RNA-protein complexes were predicted poorly, the rest of the targets were predicted with high accuracy and with good topology.

The recent assessment report mentions that several predictor groups (17/40 participants) utilized deep learning methods, including DeepFoldRNA [2] and E2Efold-3D [5], for predicting the 3D structure of the targets. None of the predictors that used deep learning for RNA 3D structure prediction took one of the top three positions, which was assigned by the authors to the fact that there is a small amount of high-resolution experimentally determined RNA structures available to train the deep learning models.

### DeepFoldRNA and trRosettaRNA

DeepFoldRNA [2] and trRosettaRNA [3] are novel deep-learning applications for sequence-based RNA 3D structure prediction. Both algorithms are inspired by AlphaFold [22] and trRosetta [23] that have recently accurately predicted the 3D structure of proteins. DeepFoldRNA and trRosettaRNA consist of a geometry prediction component and a 3D folding component based on energy minimization. Firstly, the geometric restraints are predicted by a transformer network and secondly they are used as restraints to guide energy minimization by the L-BFGS algorithm.

DeepFoldRNA and trRosettaRNA deploy a 48-block transformer network for predicting geometries. The transformer network receives as input the multiple sequence alignment (MSA) representation and the secondary structure representation and with the mechanism of self-attention updates the two representations, effectively highlighting important features and relationships that are relevant to the accurate prediction of the geometries.

The main difference in methodology followed by DeepFoldRNA and trRosettaRNA lies in the 3D folding procedure. DeepFoldRNA includes only 4 atoms of each nucleotide (P, C4’, N1 for pyrimidines and N9 for purines and C2 for adenine and uracil, C4 for cytosine, C6 for guanine) in energy minimization and then adds the rest of atoms in the 3D structure through refinement, using SimRNA [24] and QRNAS [25]. On the other hand, trRosettaRNA includes all atoms in energy minimization. The authors of the trRosettaRNA paper claim that this full-atom optimization approach leads to better accuracy in predicting the side chains of RNA compared to DeepFoldRNA [2].

### RhoFold (E2Efold-3D)

RhoFold, alternatively named as E2Efold-3D, is a deep neural network for *de novo* RNA 3D structure prediction [5]. RhoFold receives as input an MSA and a pairwise representation that it passes through three consecutive modules: the pre-trained RNA Foundation Module for enriching the MSA, the E2Eformer Module for feature extraction from the enriched MSA and the pairwise representation and finally, the Structure Module that predicts the positions of the base frames in the structure. The base frames are defined as the (C4′, C1′, N1/N9) triangles. Inside the Structure Module, the Invariant Point Attention (IPA) submodule predicts the rotation and translation matrices of each base frame, starting from the MSA and pairwise representation resulted from the E2Eformer Module and calculates the full-atoms coordinates. The loss function of RhoFold penalizes the discrepancies in the 3D structure (based on clashscore, plDDT (ref.), FAPE (ref.) and P, C4, N atoms distances in all pairs of residues), the 2D structure (predicted base-pair - non-base-pair matrix) and the base-pairs distances formed in the predicted 3D structure. It also penalizes based on the residues predicted to covary in the MSA, with a language model masking approach.

RhoFold is a third algorithm for RNA 3D structure prediction, that for predicted 3D candidate models for the validation and testing datasets that we used for developing and evaluating our method. RhoFold finishes with the refinement of the predicted 3D structure using the AMBER force field relaxation for 1000 cycles. We included both the unrelaxed and relaxed 3D structure in our datasets.

RhoFold shares architectural similarities with DeepFoldRNA and trRosettaRNA in terms of feature extraction from MSA and pairwise representations, but it diverges in the methodology used to predict the final RNA 3D structures. RhoFold opts for an end-to-end trainable approach using IPA, whereas DeepFoldRNA and trRosettaRNA rely on energy minimization techniques with constraints informed by their respective neural network predictions.

### Atomic Rotational Equivariant Scorer (ARES)

The Atomic Rotational Equivariant Scorer (ARES) [4] is a deep neural network that predicts the RMSD of a 3D structure from the experimentally determined 3D structure by the coordinates of its atoms. ARES uses 3D equivariant convolution [26] in order to capture the geometric patterns in the 3D structure, at different positions and orientations. The atoms of the structure pass through 3 layers of 3D equivariant convolution, progressively capturing broader structural elements in the structure. To predict the accuracy of different RNA 3D structure models, ARES was integrated with FARFAR2 [27] another computational method for constructing RNA 3D structures from nucleotide sequence. ARES was trained on 14 RNAs, with lengths ranging from 17 to 47 nucleotides, including one pseudoknot (PDB: 1L2X). For these RNAs, 1000 FARFAR2 models were used from the FARFAR2-Classics collection of models.

During the RNA-Puzzles IV round, ARES successfully identified the most accurate models from the candidate FARFAR2 models for four targets, placing the ARES team at the top of the RMSD scoring table for those specific RNAs [4]. Additionally, in a separate test set of 16 RNAs, each with 5000 FARFAR2 models, ARES achieved a median RMSD of approximately 11Å. This performance significantly outperformed older scoring methods in terms of median RMSD estimation [4].

### Molprobity

Molprobity is a web service for the validation of the structures of proteins, nucleic acids and protein-nucleic acids complexes [28]–[31]. Molprobity evaluates the geometry of the structures in a spatially restricted environment, assessing factors such as steric clashes, bond lengths and angles. The Molprobity summary table provides the total number and the percentage (or distribution percentile) of the stereochemical discrepancies over the whole structure which can be used for the evaluation of the structure. For RNA 3D structures, the statistics that are included in the Molprobity summary are the following:

1. Clashscore: the number of serious steric clashes (clash of Van der Waals surfaces between 2 atoms > 0.4 Å) per 1000 atoms in the structure [30], [31].
2. Bad angles (%): the percentage of nucleotides in the structure with backbone angle smaller or larger than at least 4σ from the ideal value [30].
3. Bad bonds (%): the percentage of nucleotides in the structure with backbone bonds with length smaller or larger than at least 4σ from the ideal value [30].
4. Bad backbone conformations (%): the percentage of RNA suites (sugar-to-sugar units in the RNA backbone) in the structure that are not known rotamers [30].
5. Probably wrong sugar puckers (%): the percentage of sugar puckers in the structure that have been characterized as C3’-endo while being C2’-endo [30].
6. Chiral handedness swaps (%): the percentage of chiral centers in the structure with distortions in chiral volume (> 4σ) that lead to chiral handedness swaps [29].
7. Tetrahedral geometry outliers (%): the percentage of tetrahedral centers with distortions in chiral volume that are large, but don’t cause chiral handedness swaps [29].

For the training set used in our work we utilized DeepFoldRNA and trRosettaRNA for predicting the 3D structure of RNAs with pseudoknots. On those predicted structures we trained an artificial neural network called REA (RMSD Estimation Algorithm) to predict the root mean squared deviation (RMSD) of the predicted structures to the experimentally determined structure using their Molprobity statistics. Our algorithm aims to facilitate the evaluation and selection of the most accurate among the candidate RNA 3D structure models.

## Results

### RMSD Estimation Algorithm (REA)

We have built the RMSD Estimation Algorithm (REA), a feed-forward neural network with one hidden layer. REA receives as input the Molprobity features of a 3D RNA structure and predicts its RMSD to its experimentally determined structure. REA was trained on 182 3D RNA structures listed in Supp. Table 3 predicted by DeepFoldRNA and trRosettaRNA (DeepFoldRNA predicts six alternative structures for each sequence while trRosettaRNA predicts one. 26 sequences in the training set give (6+1)*26=182 predicted structures). The RNA sequences that were used as the starting point for the 3D RNA structure predictions belonged to pseudoknotted RNAs. The 3D structure models in the training set corresponded to 26 RNAs with pseudoknots (Supplementary, Table 1). REA was tested on a test set of 69 models of RNAs that form pseudoknots (Supplementary, Table 2) and on another test set of 117 models of RNAs that don’t form pseudoknots (Supplementary, Table 3). The development of the model was performed with 10-fold cross-validation on the training set.

### Hyperparameter tuning of REA

Hyperparameter tuning aimed to improve the predictive performance or generalization ability of the RMSD prediction of a 3D RNA structure from Molprobity features. Initially, the TPE Sampler from Optuna [32] was employed to optimize the activation function, learning rate, batch size, maximum number of epochs, and number of hidden units. The hyperparameter ranges that were provided to the TPE sampler were the same as the ones that have resulted in optimum performance for the initial energies-only model. The sum of the mean squared error in the training (MSE train) and validation set (MSE validation) was set as the objective function to be minimized by the TPE Sampler, as set in the initial energies-only model. The optimal hyperparameter values that emerged from the search with TPE sampler for the hyperparameters of REA were: 10 number of hidden units, learning rate 0.001, ReLU activation function, maximum number of epochs 150 and batch size 32.

Next, we focused on refining specific hyperparameters while keeping the activation function (ReLU) and batch size (32) fixed. We explored various combinations of learning rates, number of hidden units, maximum number of epochs, and feature sets with cross-validation. Learning rates of 0.001, 0.0002, 0.0001, and 0.00005 were tested, along with 5, 10, and 15 hidden units, and maximum number of epochs of 150 and 300. Two feature sets were considered: Molprobity features only and Molprobity features + Energies. Interestingly, it was observed that the Molprobity + Energies feature set yielded inferior results, indicating a potential trade-off between the two sets.

Finally, a more extensive search was conducted to refine the learning rate and number of hidden units, aiming to optimize these critical hyperparameters further.

### Combinations of hyperparameters and comparison between the Molprobity and Molprobity + Energies feature sets

Starting from the learning rate 0.001, the number of hidden units 10 and the maximum number of epochs 150 found by the TPE Sampler, we searched further for optimal values for these hyperparameters in combination. Learning rates of 0.001, 0.0002, 0.0001, and 0.00005 were tested, along with 5, 10, and 15 hidden units, and maximum number of epochs of 150 and 300.

We also tuned the features set in combination with the learning rate, the number of hidden units and the maximum epochs. We tested the Molprobity and the Molprobity+Energies features sets. The Molprobity+Energies includes the Molprobity features for the RNA 3D structure and the energies calculated by DeepFoldRNA during folding and structure refinement. In particular, in the Molprobity+Energies set the energies are: the energy after refinement (te), the energy after folding without refinement (tewor) and their difference (te-tewor).

Each combination of hyperparameters was evaluated using a 10-fold cross-validation approach with 5 iterations per data split. The mean squared error on the validation set (MSE CV) was used as the evaluation metric during cross-validation. The best performing model during cross-validation was determined by selecting the model with the minimum MSE on the validation set. We used only the DeepFoldRNA models from the training set (156 structures) in the cross-validation experiments, because they contain the required energy related features.

Figure 1, Figure 2, and Figure 3 present the learning curves of the best performing model from each cross-validation cycle for three different hyperparameter combinations. Figure 1 displays the learning curves for the hyperparameter combination of learning rate 0.001, number of hidden units 10, and max epochs 150. Figure 2 presents the learning curves for the hyperparameter combination of learning rate 0.0001, number of hidden units 10, and max epochs 300. Lastly, Figure 3 shows the learning curves for the hyperparameter combination of learning rate 0.001, number of hidden units 15, and max epochs 300. By analyzing these learning curves, we can evaluate the performance and generalization ability of the model during the training process. This analysis enables us to select the hyperparameter combination that yields the best performance.

**Figure 1:**
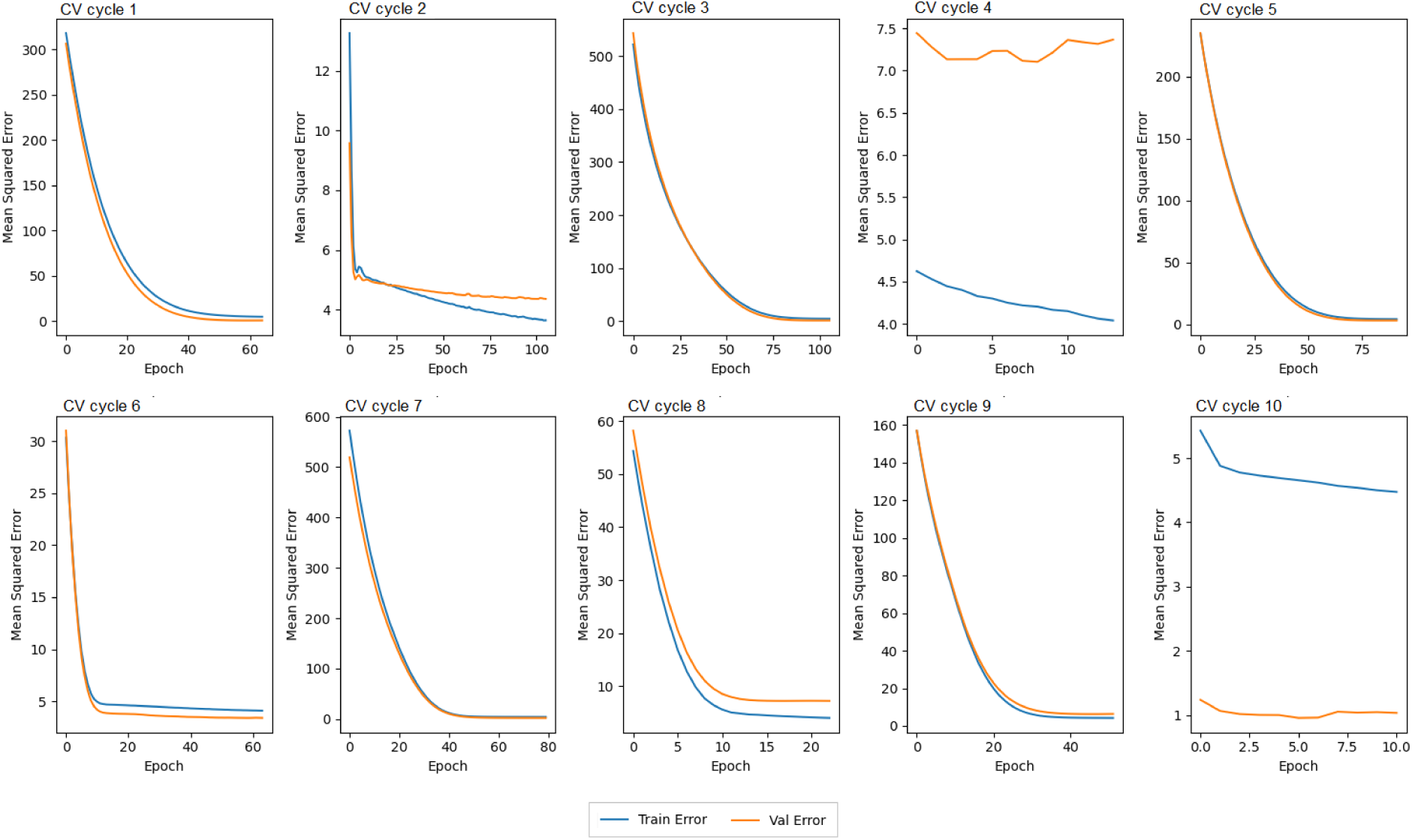
The learning curves of the best performing model of every cross-validation cycle for the hyperparameters 0.001 learning rate, 10 number of hidden units and 150 maximum number of epochs. This combination of hyperparameters represents the initial hyperparameter values obtained from the TPE Sampler. The models were trained using early stopping of 5 epochs.

**Figure 2:**
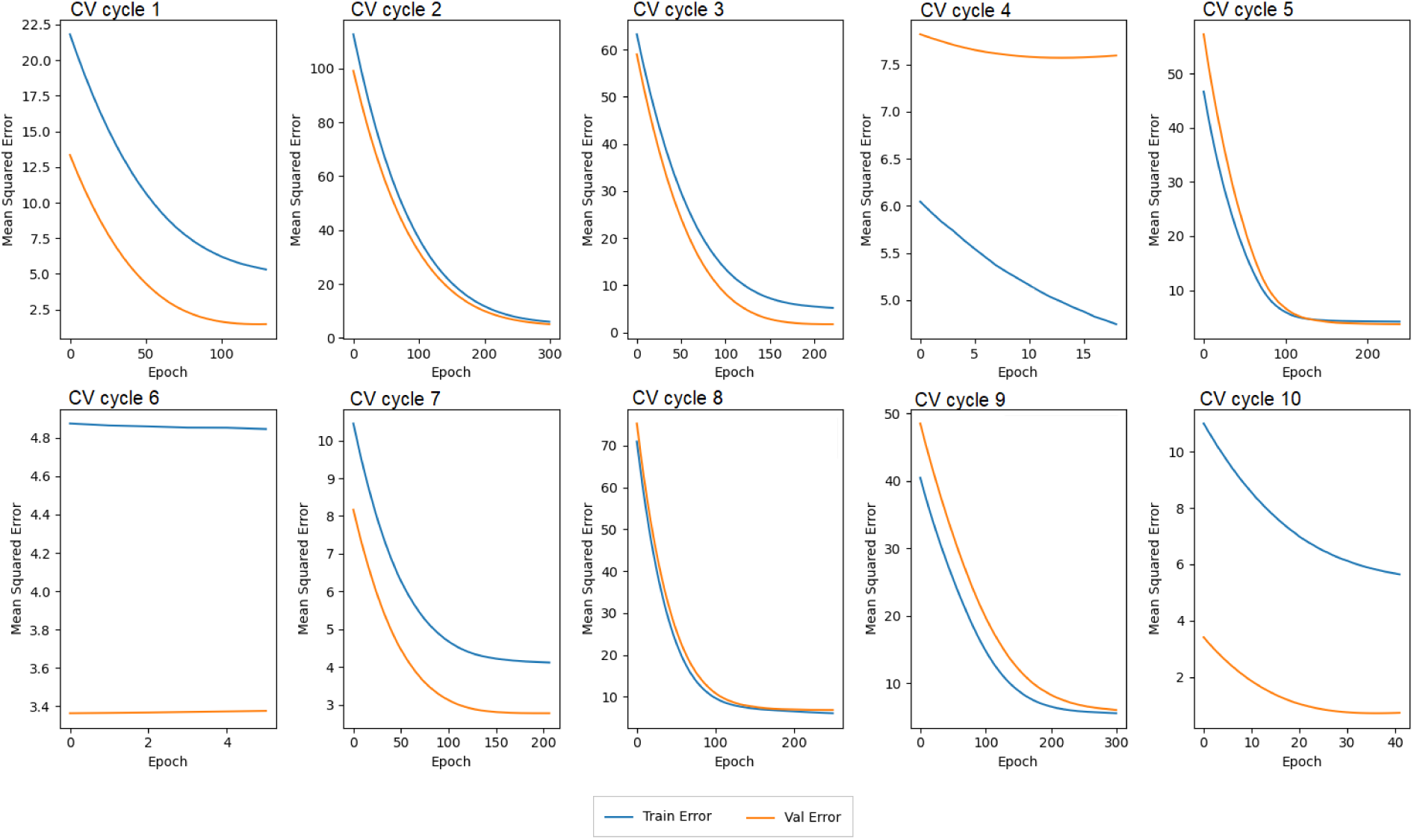
The learning curves of the best performing model of every cross-validation cycle for the hyperparameters 0.0001 learning rate, 10 number of hidden units and 300 maximum number of epochs. In this combination of hyperparameters the learning rate is reduced by an order of magnitude, resulting in slower convergence. The models were trained using early stopping of 5 epochs.

**Figure 3:**
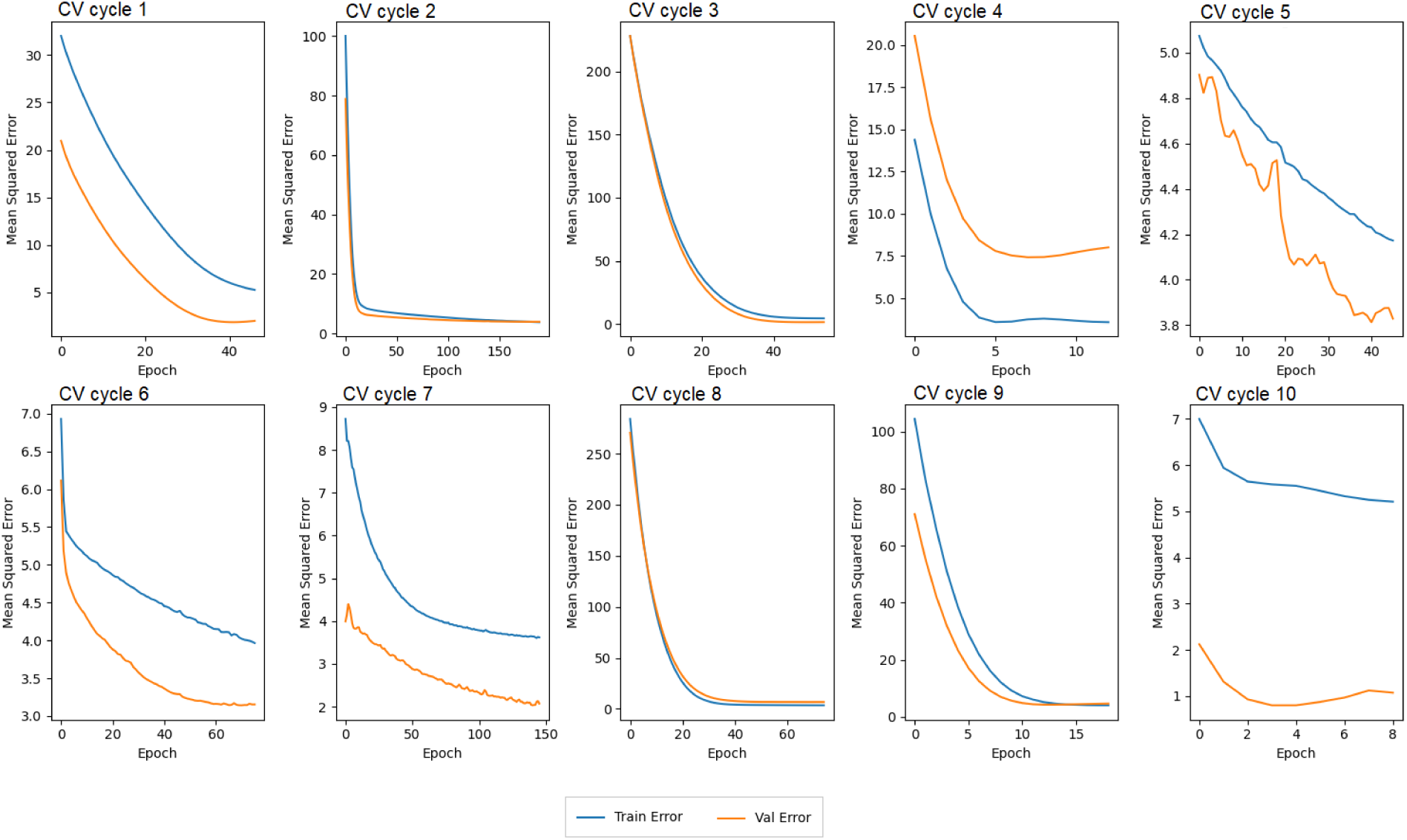
The learning curves of the best performing model of every cross-validation cycle for the hyperparameters 0.001 learning rate, 15 number of hidden units and 300 maximum number of epochs. This hyperparameters combination features an increased number of hidden units from 10 to 15, with both the training MSE and validation MSE decreasing in parallel for most models, indicating effective learning. The models were trained using early stopping of 5 epochs.

The results shown in Figure 1, Figure 2 and Figure 3 indicate a direction towards the optimal combination of learning rate and number of hidden units. Among the three combinations investigated, the combination of learning rate 0.001 and 15 number of hidden units shows fast convergence and good generalization ability.

Figure 4, Figure 5 and Figure 6 show the performance of the model during cross-validation with respect to the learning rate, the number of hidden units and the maximum number of epochs respectively, for the two feature sets.

**Figure 4:**
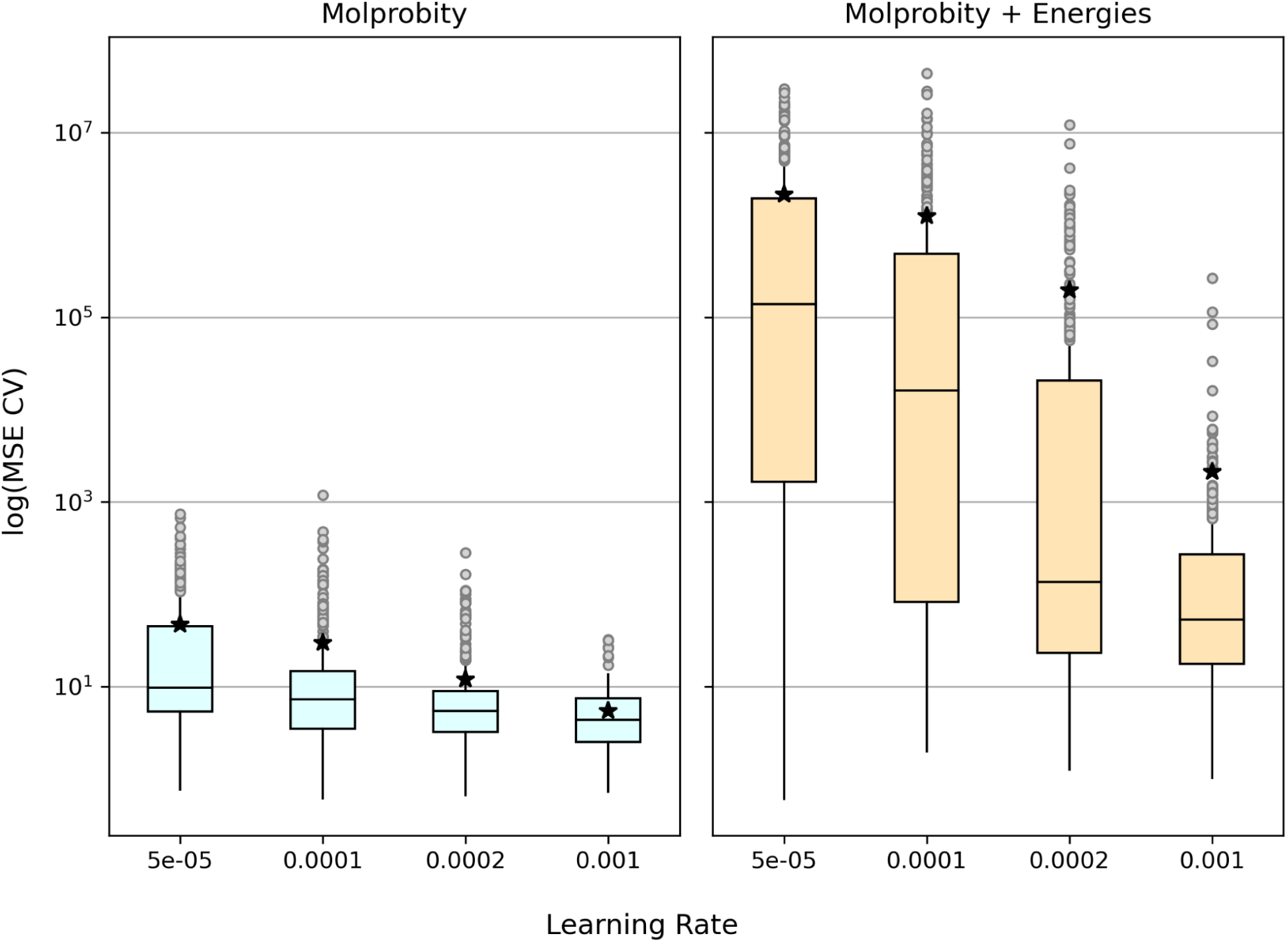
Comparison of cross-validation MSE (logarithmic scale) for different learning rates between Molprobity and Molprobity + Energies feature sets. The average of every distribution is shown with a star.

**Figure 5:**
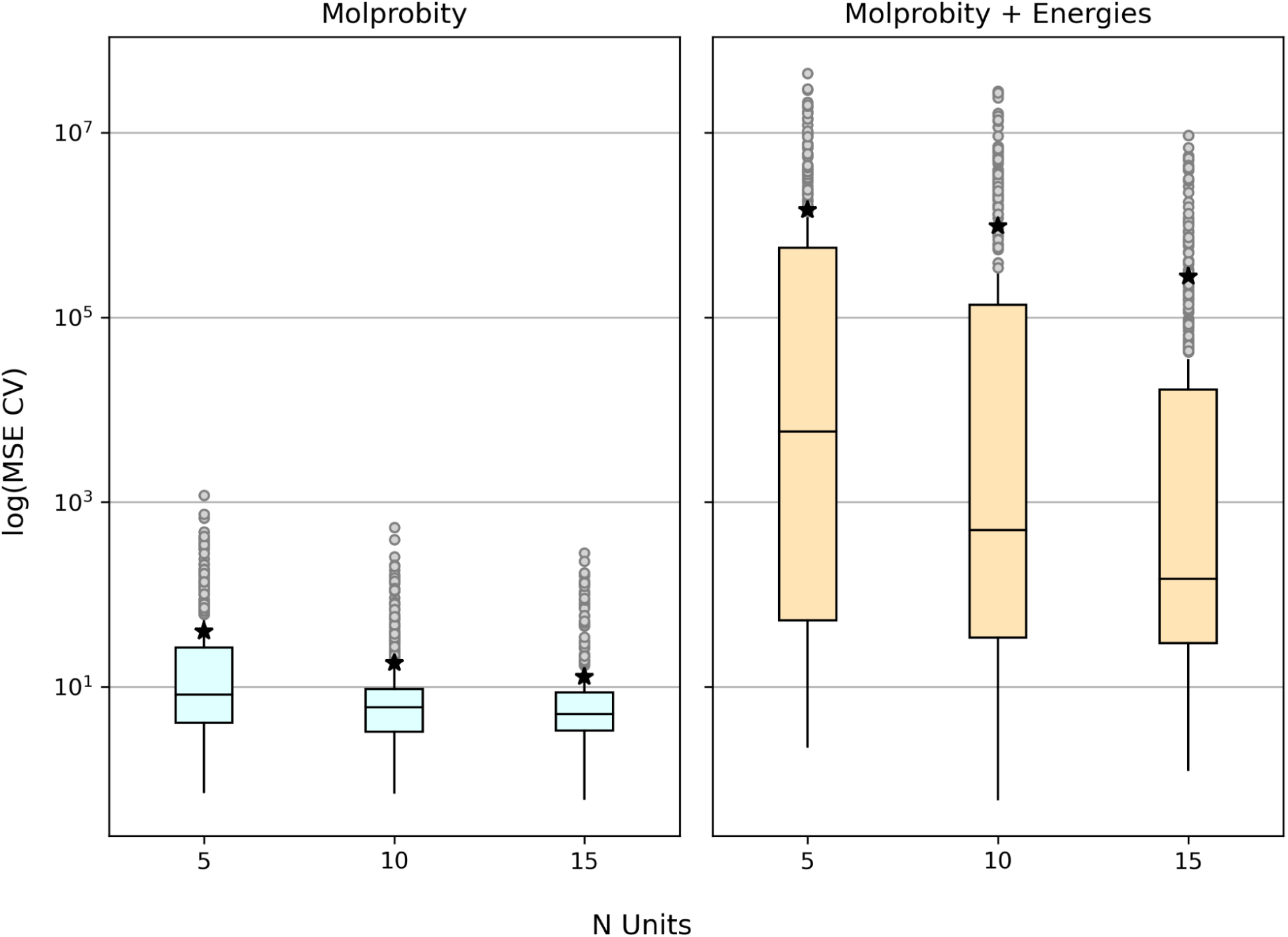
The logarithm of cross-validation MSE for different numbers of hidden units separated between Molprobity and Molprobity + Energies features sets.

**Figure 6:**
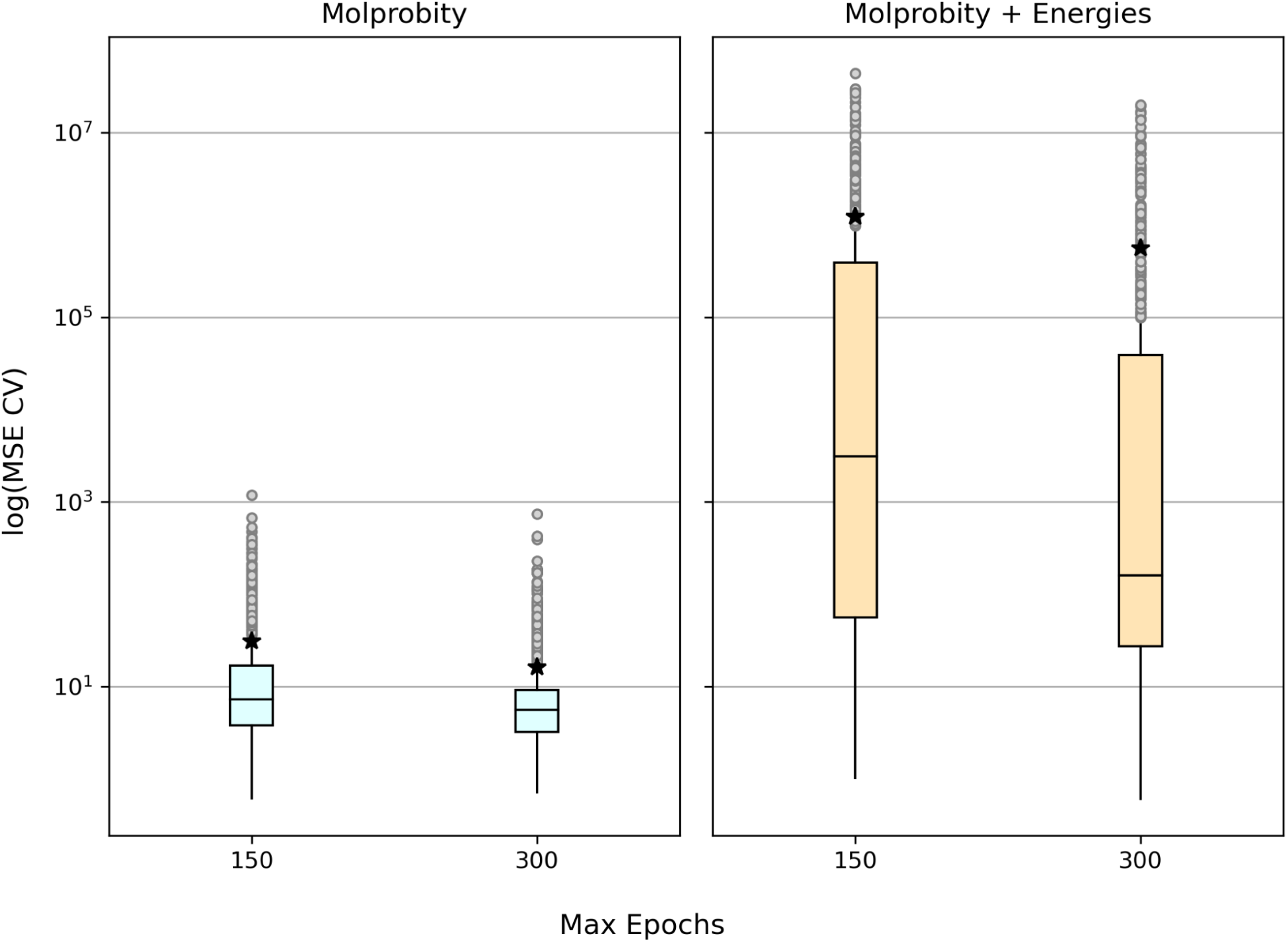
The logarithm of cross-validation MSE for different numbers of maximum epochs separated between Molprobity and Molprobity + Energies features sets.

Figure 4 presents the performance of the model during cross-validation using different learning rates for the Molprobity and Molprobity + Energies feature sets. The learning rates compared are 0.00005, 0.0001, 0.0002, and 0.001. Upon examining the pair-wise comparison of the same learning rate between the two feature sets, notable differences in the log(CV MSE) distributions are observed, with differences spanning at least one order of magnitude. The performance using the Molprobity feature set is superior to that of the Molprobity + Energies feature set. Focusing on the Molprobity feature set, we observe that a learning rate of 0.001 yields the smallest median and interquartile range (IQR) of log(CV MSE) values (Supplementary, Table 5). This indicates that, among the evaluated learning rates, 0.001 leads to the best performance when using the Molprobity feature set of the DeepFoldRNA structures.

Figure 5 presents the performance of the model during cross-validation using different numbers of hidden units for the Molprobity and Molprobity + Energies feature sets. The numbers of hidden units compared are 5, 10 and 15. Upon examining the pair-wise comparison of the same number of hidden units between the two feature sets, notable differences in the log(CV MSE) distributions are observed, with differences spanning at least one order of magnitude. As it was noticed for the different learning rates, the performance using the Molprobity feature set is superior to that of the Molprobity + Energies feature set. Focusing on the Molprobity feature set, it is observed that 15 hidden units yield the smallest median and IQR of log(CV MSE) values (Supplementary, Table 6). This indicates that, among the evaluated numbers of hidden units, 15 produces the best performance when using the Molprobity feature set of the DeepFoldRNA structures.

Figure 6 presents the performance of the model during cross-validation using different numbers of maximum epochs for the Molprobity and Molprobity + Energies feature sets. The numbers of maximum epochs compared are 150 and 300. Upon examining the pair-wise comparison of the same number of maximum epochs between the two feature sets, notable differences in the log(CV MSE) distributions are observed, with differences spanning at least one order of magnitude. As it was noticed for the different learning rates and different numbers of hidden units, the performance using the Molprobity feature set is superior to that of the Molprobity + Energies feature set. Focusing on the Molprobity feature set, it is observed that 300 maximum epochs yield the smallest median and IQR of log(CV MSE) values (Supplementary, Table 7). This indicates that, between 150 and 300 number of maximum epochs, 300 produces the best performance when using the Molprobity feature set of the DeepFoldRNA structures.

Based on the above cross-validation experiments that compare different combinations of learning rates, number of hidden units, maximum number of epochs and feature sets, using DeepFoldRNA models, we select the Molprobity features set for REA. We also select 300 as the maximum number of training epochs. We also have indications that a learning rate around 0.001 or higher and a number of hidden units around 15 or higher are going to be optimal for our task.

### Refinement of learning rate and number of hidden units

The learning rate and the number of hidden units are critical hyperparameters. Finding an appropriate learning rate is crucial for achieving good performance and efficient convergence. Finding an appropriate number of hidden units equals the balance between model complexity and generalization capacity. The refinement of the learning rate and the number of hidden units involved searching a more extensive set of values for each.

Each hyperparameter value was evaluated using a 10-fold cross-validation approach with 5 iterations per data split. The mean squared error (MSE) on the validation set was used as the evaluation metric during cross-validation. We used all the models from the training set (182 structures) in the cross-validation experiments, DeepFoldRNA and trRosettaRNA models included.

We searched for the optimal learning rate in the range 5e-5 to 1.28e-2 in steps, by adding half of the value of the previous step to the next step. Overall, we tested the values 0.00005, 0.0001, 0.0002, 0.0004, 0.0008, 0.0012, 0.0016, 0.0024, 0.0032, 0.0048, 0.0064, 0.0096, 0.0128. Figure 7 shows the cross-validation performance of the different learning rates.

**Figure 7:**
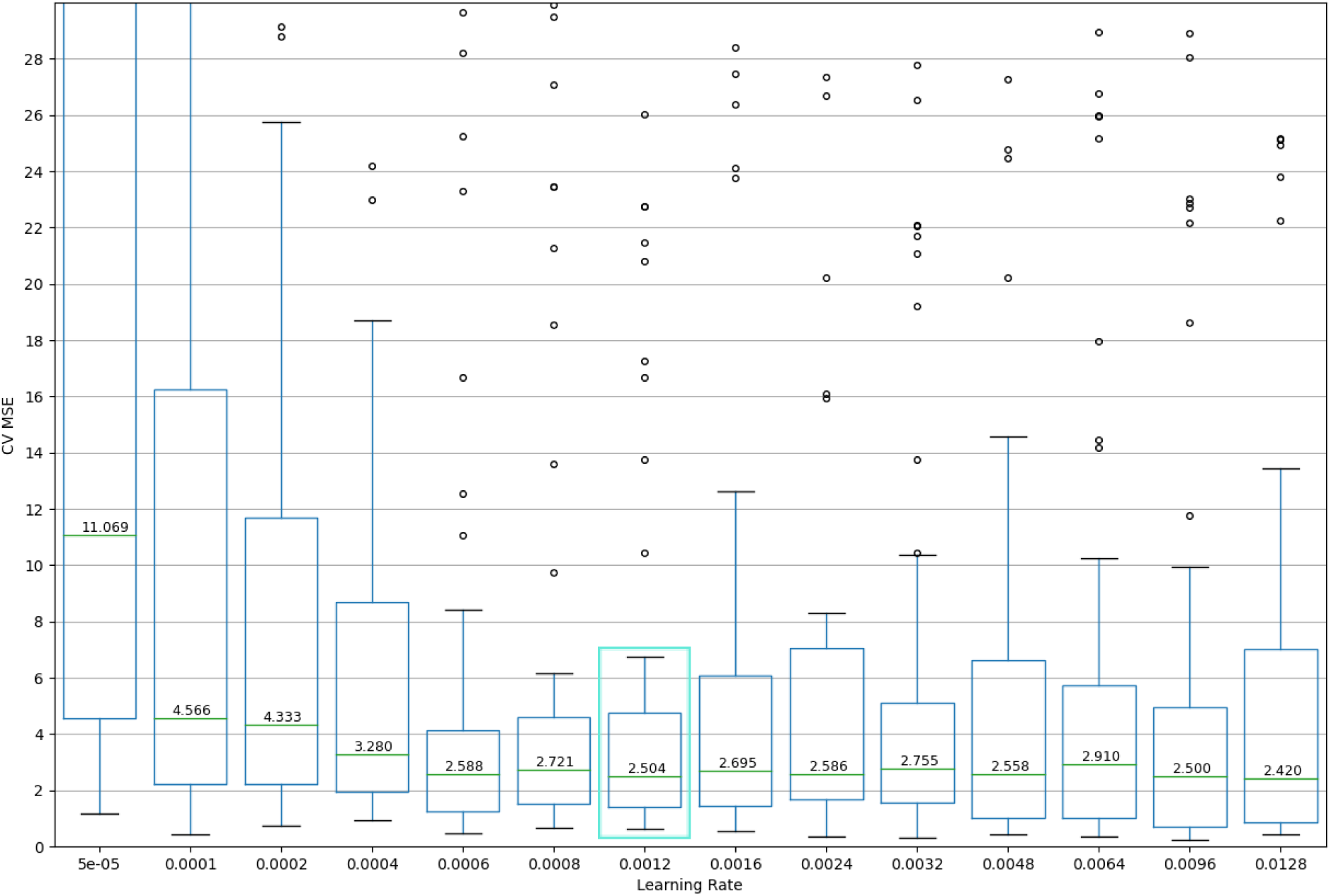
Cross-validation MSE for different learning rates while keeping other hyperparameters fixed (hidden units = 10, max epochs = 300, batch size = 32). Each box plot represents the MSE distribution from a 10-fold cross-validation with 5 iterations. The learning rate 0.0012 (marked with green rectangle) demonstrates a combination of low median CV MSE and narrow spread.

In Figure 7, the comparison among the different learning rates distinguishes 0.0012 as the optimal learning rate with the third smallest CV MSE median (2.504) and narrow IQR (3.35). Although the learning rate of 0.0128 has the smallest median CV MSE (2.420), its IQR of CV MSE values (6.16) is wider. Similarly, the learning rate of 0.00096 achieves the second smallest CV MSE (2.500), but its IQR (4.23) is wider compared to 0.0012. Based on this evaluation, it appears that the learning rate of 0.0012 demonstrates a desirable combination of low CV MSE median and a narrow IQR of MSE values (Supplementary, Table 8), indicating better performance and consistency across the cross-validation folds.

After having selected 0.0012 learning rate, we proceeded in tuning the number of hidden units. We tested all numbers from 1 to 20 with a step of 1 and from 20 to 60 with a step of 20. Figure 8 shows the cross-validation performance of the different numbers of hidden units.

**Figure 8:**
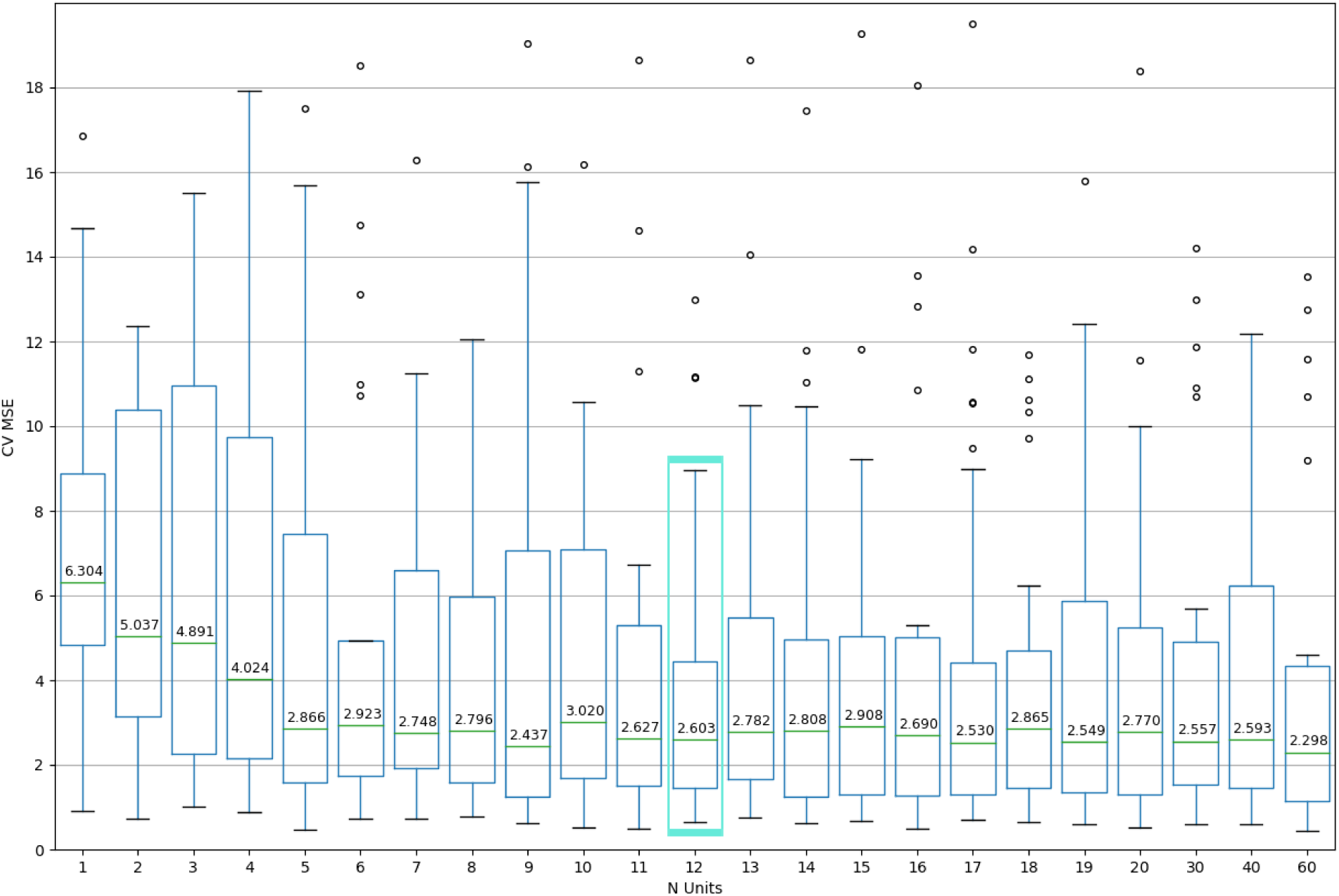
Cross-validation MSE for different number of hidden units while keeping other hyperparameters fixed (learning rate = 0.0012, max epochs = 300, batch size = 32). Each box plot represents the MSE distribution from a 10-fold cross-validation with 5 iterations. Twelve (12) is considered to be the optimal number of hidden units, because of its small MSE CV and small IQR during cross-validation, which indicates consistency in performance.

In Figure 8, the comparison among the different numbers of hidden units distinguishes 12 as the optimal number of hidden units with a small CV MSE median (2.60) and narrow IQR (3.00). The smallest median (2.30) belongs to 60 hidden units, which on the contrary shows a larger IQR (3.20) (Supplementary, Table 9). Additionally, the choice of 12 hidden units keeps the complexity and training time of the model as low as possible.

After having selected 12 as the optimal number of hidden units we tune the learning rate for one last time, in order to search for an optimal number of hidden units and learning rate combination. Figure 9 shows the cross-validation performance of the different learning rates during the second refinement.

**Figure 9:**
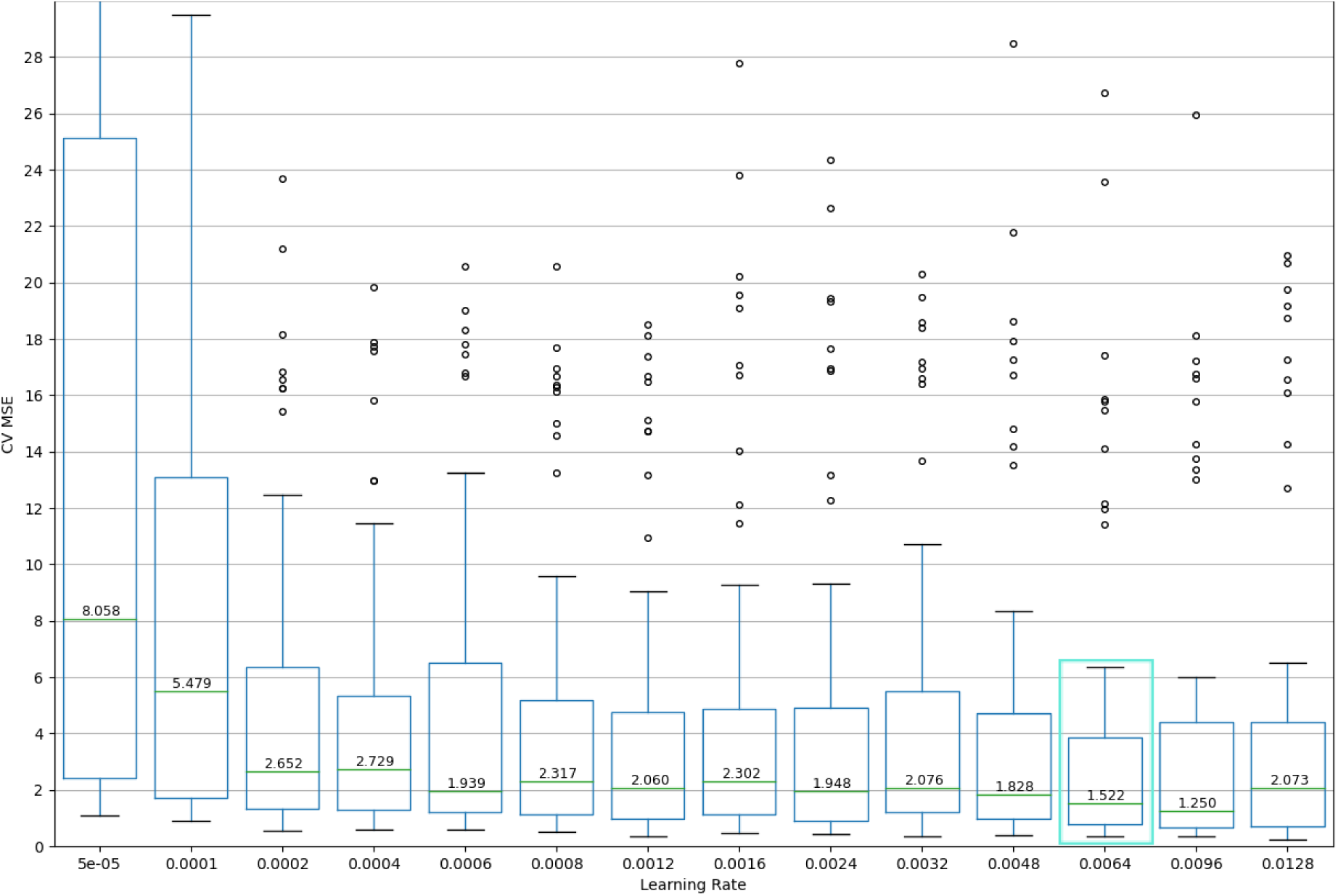
Cross-validation MSE for different learning rates, after determining the number of hidden units (12) and while keeping the other hyperparameters fixed (max epochs = 300 and batch size = 32). Each box plot represents the MSE distribution from a 10-fold cross-validation with 5 iterations. The learning rate of 0.0064 emerges as the new optimal choice.

In Figure 9, the comparison of performance during cross-validation among the different learning rates distinguishes 0.0064 as the optimal learning rate with the second smallest CV MSE median (1.52) and the narrowest IQR (Supplementary, Table 10). The second refinement led to improvement of ∼1 unit in the CV MSE, from 2.50 after the first refinement to 1.52 after the second refinement. Starting with a performance of median MSE CV 4.39 by the initial model with hyperparameters proposed by TPE Sampler, after further hyperparameter tuning we achieved a decrease of almost 3 units with final MSE CV 1.52.

### Feature selection

The selection of features aims at improving the performance of the model, assessing feature importance and reducing the computational cost of modeling. Feature selection improves the performance of the model by selecting among the Molprobity features those that are the most relevant to the RMSD prediction and demonstrate small correlation to each other. Feature selection ensures that the chosen features provide valuable information without redundancy, enabling the network to focus on the specific features that significantly contribute to accurate RMSD prediction. Additionally, given the small number of training examples the reduction of the data dimensionality can reduce the number of parameters in the neural network, prevent overfitting and improve performance.

We enhanced the Molprobity feature set with two more features: “length_target_norm”, the normalized length of RNA, and the “Clashscore all atoms percentile”, calculated by Molprobity is the percentile of the “Clashscore all atoms” distribution of structures with similar resolution that the “Clashscore all atoms” score belongs to. We added the “length_target_norm” to let the model identify patterns related to the length of RNA molecules and their structural characteristics. Additionally, we added the “Clashscore all atoms percentile” feature to include information about the severity of the clashes in the structure that is shown through this relative measure. The addition of “length_target_norm” and “Clashscore all atoms percentile” features reduced the median MSE CV from 5.02 to 3.71 (Supplementary, comparison between Tables 11 and 12, learning rate 0.0064) and improved the performance of the model.

We normalized the length of the target RNAs in the training set, before the train/validation split. We normalized by dividing with the maximum length, which the majority of splits was in the training set and not in the validation. Also we performed the train/validation split so that the median of length of RNAs in the total training set to be equal (± length std. error) from the median length in the validation set. The similarity in the distribution of the length in training and validation set along with the further evaluation of the model in test sets show that we can trust the generalization ability shown by the model.

We performed knockout experiments for selecting the Molprobity features that lead to optimal performance. We excluded the Molprobity features one by one and then in pairs from the features and we evaluated the performance of the model using the remaining features. The models using different subsets of the Molprobity features were evaluated using cross-validation. Each subset of features was evaluated using a 10-fold cross-validation approach with 20 iterations per data split. The mean squared error (MSE) on the validation set was used as the evaluation metric during cross-validation. Figure 10 shows the performance of models with different features knocked out during cross-validation.

**Figure 10:**
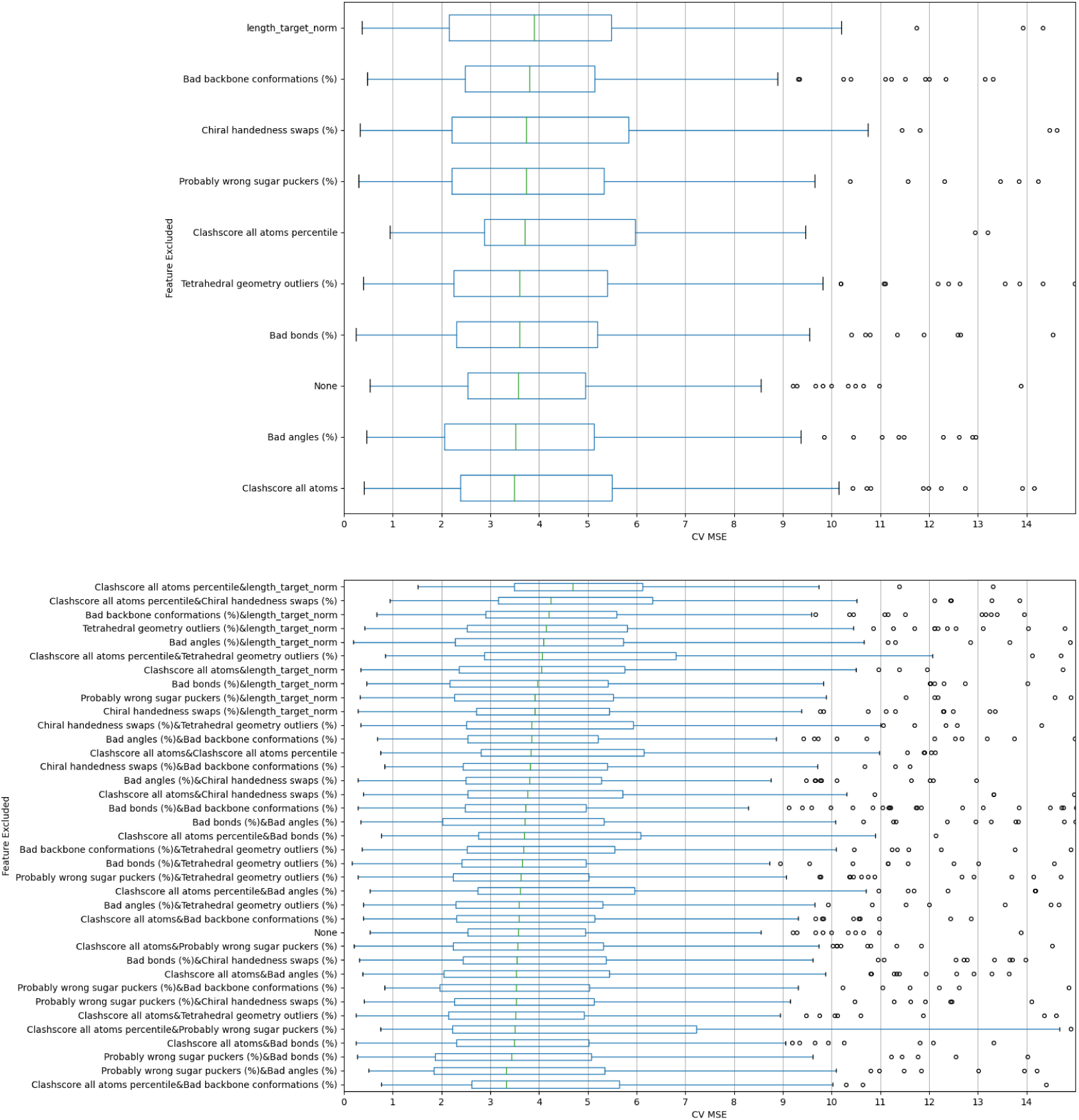
Cross-validation (10-fold with 20 iterations) performance of Molprobity features sets where different features were knocked out. Single knockouts (upper) and double knockouts (lower). ‘None’ denotes that all Molprobity features are included. The knockouts are ordered by their effect on the median CV MSE, from the knockout with the biggest median CV MSE to the knockout with lowest median CV MSE. The features set with the lowest median CV MSE is the one with ‘Clashscore all atoms percentile’ and ‘Bad backbone conformation (%)’ knocked out.

We have moved from 5 to 20 iterations per cross-validation split in order to increase certainty about which features to exclude from the training set. We have performed the knockout experiments with 5, 10 and 20 iterations. At 5 iterations (Supplementary, Fig. 3) the combination “Clashscore all atoms Percentile & Probably wrong sugar puckers (%)” was second from the end but close to the final excluded features combination, based on median MSE CV. We moved on to 10 iterations (Supplementary, Fig. 4), where “Clashscore all atoms Percentile & Probably wrong sugar puckers (%)” was placed last. We further increased to 20 iterations to check if the “Clashscore all atoms Percentile & Probably wrong sugar puckers (%)” would still be the combination of features with the lowest median MSE CV, which was confirmed.

To ensure consistent evaluation of the performance of the model, we repeated the evaluation of different learning rates (shown in Figure 9 with 5 iterations) with 10-fold cross-validation with 20 iterations (Supplementary, Fig. 1 and Table 11). The learning rate 0.0064, which previously had the third smallest median MSE CV and then smallest IQR MSE CV in the 5-repeat cross-validation experiment, now holds the third smallest median MSE CV and the third smallest IQR MSE CV. When using the enhanced Molprobity feature set (Supplementary, Fig. 2 and Table 12), the learning rate 0.0064 holds the fourth smallest Median MSE CV and the third smallest IQR MSE CV, while the learning 0.0096 appears to be the optimal learning rate with smallest median MSE CV and the smallest IQR MSE CV. The cross-validation experiments for different learning rates performing 20 iterations and using the enhanced Molprobity feature suggest a new optimal learning rate that might need to be evaluated and tested in the future against learning rates larger than 0.0128.

Notably, the increase of the number of iterations per data split during cross-validation increases the MSE, which is an indication of accumulation of error when repeating the experiment. This accumulation of error needs to be investigated further in the future and it might take into account the initialization of the parameters of the model.

Figure 11 presents the regression plots between features and the RMSD. Figure 12 presents the regression plots between pairwise products of features and the RMSD, which depicts how interactions between features change the RMSD. In Figure 11, the feature ‘length target norm’ has, with 0.58, the highest correlation with the RMSD, which reflects the weakness of the predictors to accurately model larger targets. The other features with a higher absolute correlation are ‘Bad bonds (%)’ with 0.24 and ‘Clashscore all atoms percentile’ and ‘Bad backbone conformations (%)’ with −0.22.

**Figure 11:**
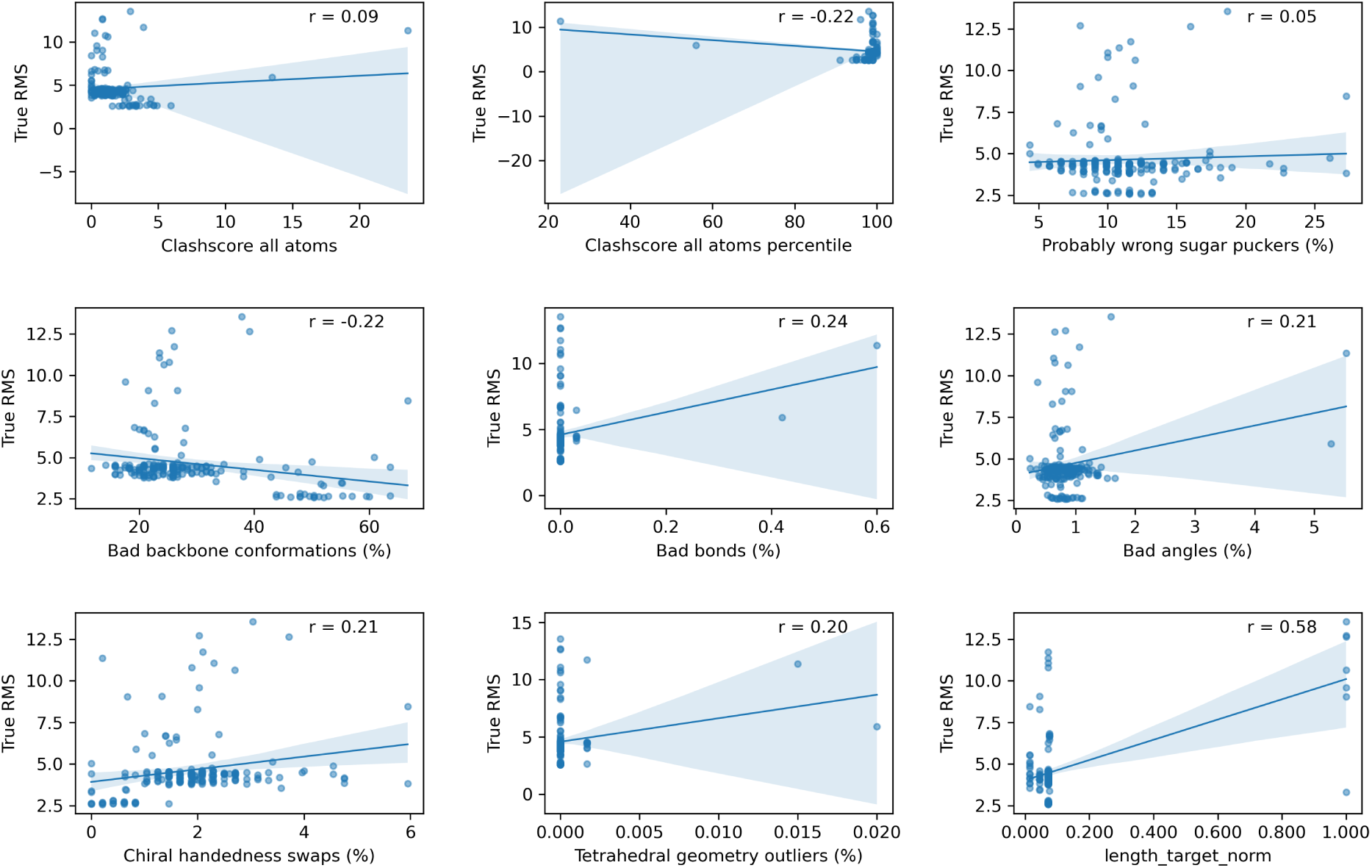
Scatter plots of RMSD against the features in the training set. The linear regression fitted line is shown in every plot, along with the Pearson’s correlation coefficient between each feature and RMSD.

**Figure 12:**
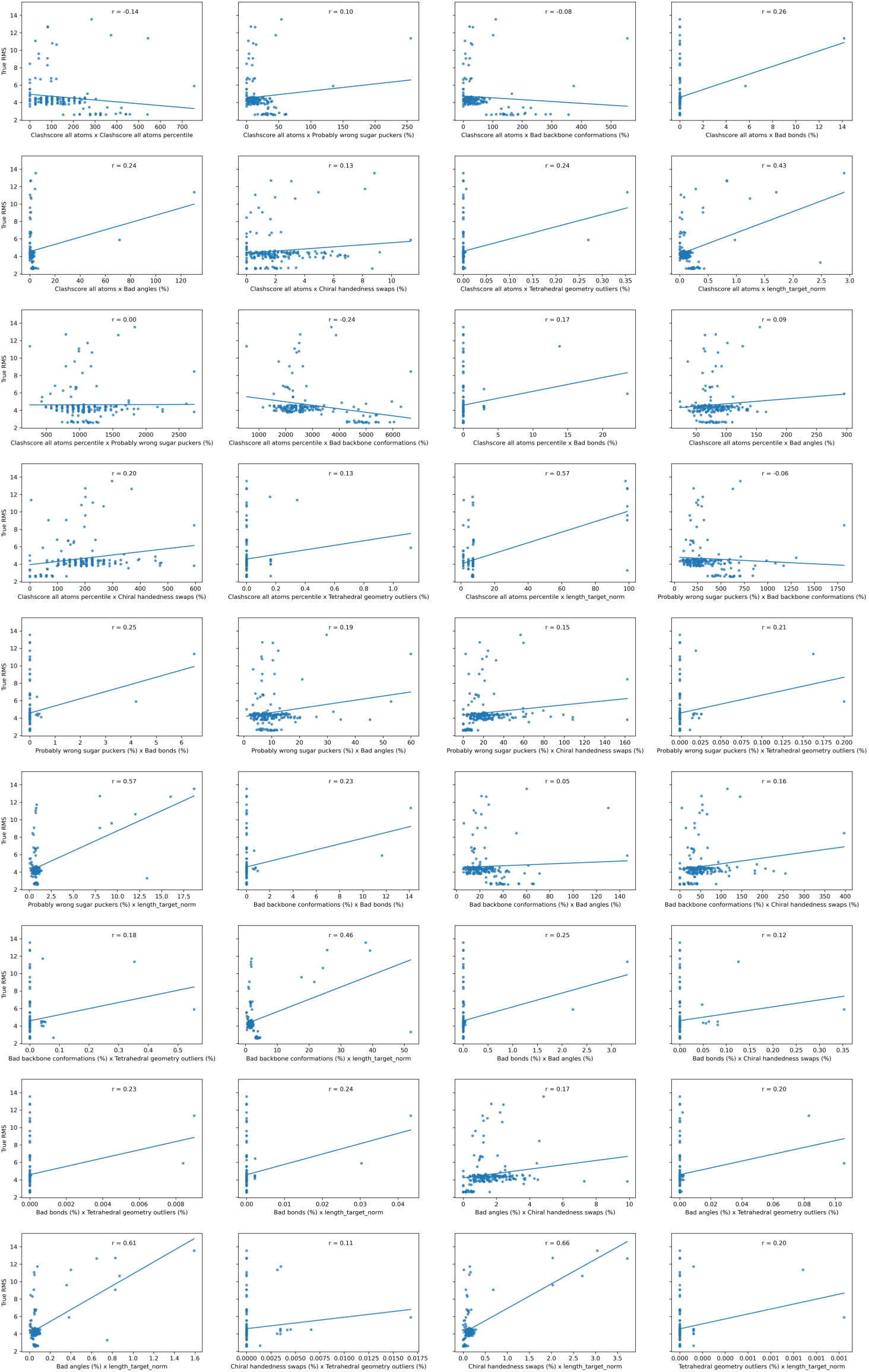
Scatter plots of RMSD against the pairwise multiplication of features in the training set. The linear regression fitted line is shown in every plot, along with the Pearson’s correlation coefficient between the products of the features and the RMSD.

Figure 10 indicates that excluding the ‘Clashscore all atoms percentile’ and the ‘Bad backbone conformations (%)’ from the features set REA achieves the lowest median MSE (3.32) during cross-validation, indicating that without these two features the model achieves the best performance. Consequently, we excluded the ‘Clashscore all atoms percentile’ and the ‘Bad backbone conformations (%)’ from the final training and testing datasets.

In Figure 12, the products with absolute correlation with RMSD greater or equal to 0.24 are considered as candidate second order features. These products are: ‘Chiral handedness swaps (%) x length target norm’ (Pearson’s correlation coefficient r = 0.66), ‘Bad angles (%) x length target norm’ (r = 0.61), ‘Probably wrong sugar puckers (%) x length target norm’ (r = 0.57), ‘Clashscore all atoms percentile x length target norm’ (r = 0.57), ‘Bad backbone conformations (%) x length target norm’ (r = 0.46), ‘Clashscore all atoms x length target norm’ (r = 0.43), ‘Clashscore all atoms x Bad bonds (%)’ (r = 0.26), ‘Probably wrong sugar puckers (%) x Bad bonds (%)’ (r = 0.25), ‘Bad bonds (%) x Bad angles (%)’ (r= 0.25), ‘Bad bonds (%) x length target norm’ (r = 0.24), ‘Clashscore all atoms x Tetrahedral geometry outliers (%)’ (r = 0.24), ‘Clashscore all atoms x Bad angles (%)’ (r = 0.24), ‘Bad bonds x length target norm’ (r = 0.24) and, ‘Clashscore all atoms percentile x Bad backbone conformations’ (r = −0.24).

In Figure 12, the products with absolute correlation with RMSD less or equal to 0.08 are considered as independent features. These features are: ‘Clashscore all atoms x Bad backbone conformations (%)’ (r = −0.08), ‘Probably wrong sugar puckers (%) x Bad backbone conformations (%)’ (r = −0.06), ‘Bad backbone conformations (%) x Bad angles (%)’ (r = 0.05) and, ‘Clashscore all atoms percentile x Probably wrong sugar puckers (%)’ (r = 0.00).

In Figure 13, the feature ‘length target norm’ is distinguished as the feature with the smallest correlation with all other features. Additionally, the higher absolute correlation is noted between the following features: ‘Clashscore all atoms percentile’ and ‘Bad bonds (%)’ (r = −0.98), ‘Clashscore all atoms’ and ‘Clashscore all atoms percentile’ (r = −0.92), ‘Bad bonds (%)’ and ‘Tetrahedral geometry outliers (%)’ (r = 0.93), ‘Bad bonds (%)’ and ‘Bad angles (%)’ (r = 0.88), ‘Bad angles (%)’ and ‘Tetrahedral geometry outliers (%)’ (r = 0.88), ‘Clashscore all atoms percentile’ and ‘Tetrahedral geometry outliers (%)’ (r = −0.88), s‘Clashscore all atoms percentile’ and ‘Bad angles (%)’ (r = −0.85), ‘Clashscore all atoms’ and ‘Bad bonds (%)’ (r = 0.85).

**Figure 13:**
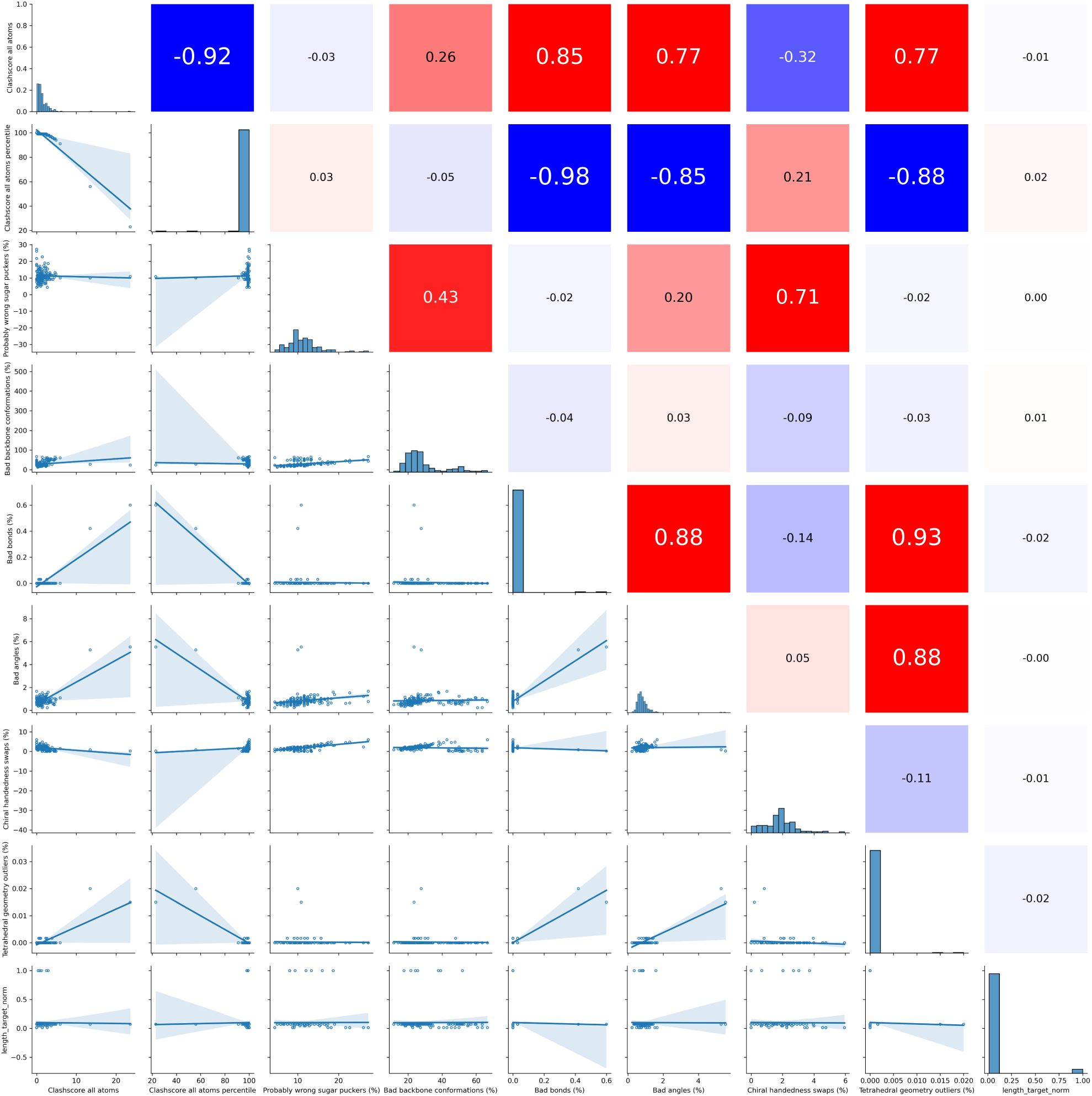
Correlation between features. (lower half) The scatter plots between all pairs of features in the training set, along with a linear regression line fit. (upper right) the Pearson’s correlation coefficients between the features pairs.

Nevertheless, the distributions of the ‘Clashscore all atoms percentile’ and ‘Bad backbone conformations (%)’ in Figure 10 and Figure 13 indicate that the multiplication of these two features results in extremely high values, shown in Figure 12. The majority of the values of ‘Clashscore all atoms percentile’ lies between 90 and 100 while the values of ‘Bad backbone conformations (%)’ span up to 60, which is high compared to the upper range of other features. Normalization of the features can help prevent the possible scenario where features with larger values dominate the modeling process and potentially mask interactions with other features.

### Support Vector Regression (SVR) model

Furthermore, we leveraged automatic machine learning methods (AutoML) [33] to identify different machine learning algorithms, apart of an artificial neural network, and feature sets that optimize the task of predicting the RMSD of an RNA structure to the experimentally determined structure using its Molprobity features. We used the JADBio platform (https://jadbio.com/) for AutoML [34], which employs Ridge Logistic Regression, Decision Trees, SVRs and Random Forests and tunes their hyperparameters. JADBio tests different methods for feature selection and selects a reference signature (or a set of equivalent signatures) for each model. We analyzed the same set used to train REA with JADBio and the model with the best performance was found to be a Support Vector Regression Machines (SVR) of type epsilon-SVR with Gaussian Kernel and hyperparameters: cost = 10.0, gamma = 1.0, epsilon = 0.1. Additionally, three signatures were proposed: 1. ‘Clashscore all atoms percentile’, ‘Chiral handedness swaps (%)’, ‘Bad backbone conformations (%)’, 2. ‘Bad angels (%)’, ‘Chiral handedness swaps (%)’, ‘Bad backbone conformations (%)’, 3. ‘Bad bonds (%)’, ‘Chiral handedness swaps (%)’, ‘Bad backbone conformations (%)’. In the beginning of the pipeline, the features were standardized.

We evaluated the three different signatures suggested by JADBio with 10-fold cross-validation with 20 iterations with leave-one-out data point per data split (Supplementary, Fig. 5 and Table 13). We kept the hyperparameters of the SVR model fixed to the proposed values. The best performing signature was signature 3 [‘Bad bonds (%)’, ‘Chiral handedness swaps (%)’, ‘Bad backbone conformations (%)’] with a MSE CV 3.51.

### Model evaluation on pseudoknotted RNAs

The evaluation of REA on pseudoknotted RNAs (Supplementary, Table 2) aims to select between REA and ARES for this specific type of complex RNA structures. We compare the RMSD given by REA and ARES using the median of the absolute difference between the predicted and true RMSD values. Figure 14 shows the distribution of the absolute differences between the predicted and the true RMSD.

**Figure 14:**
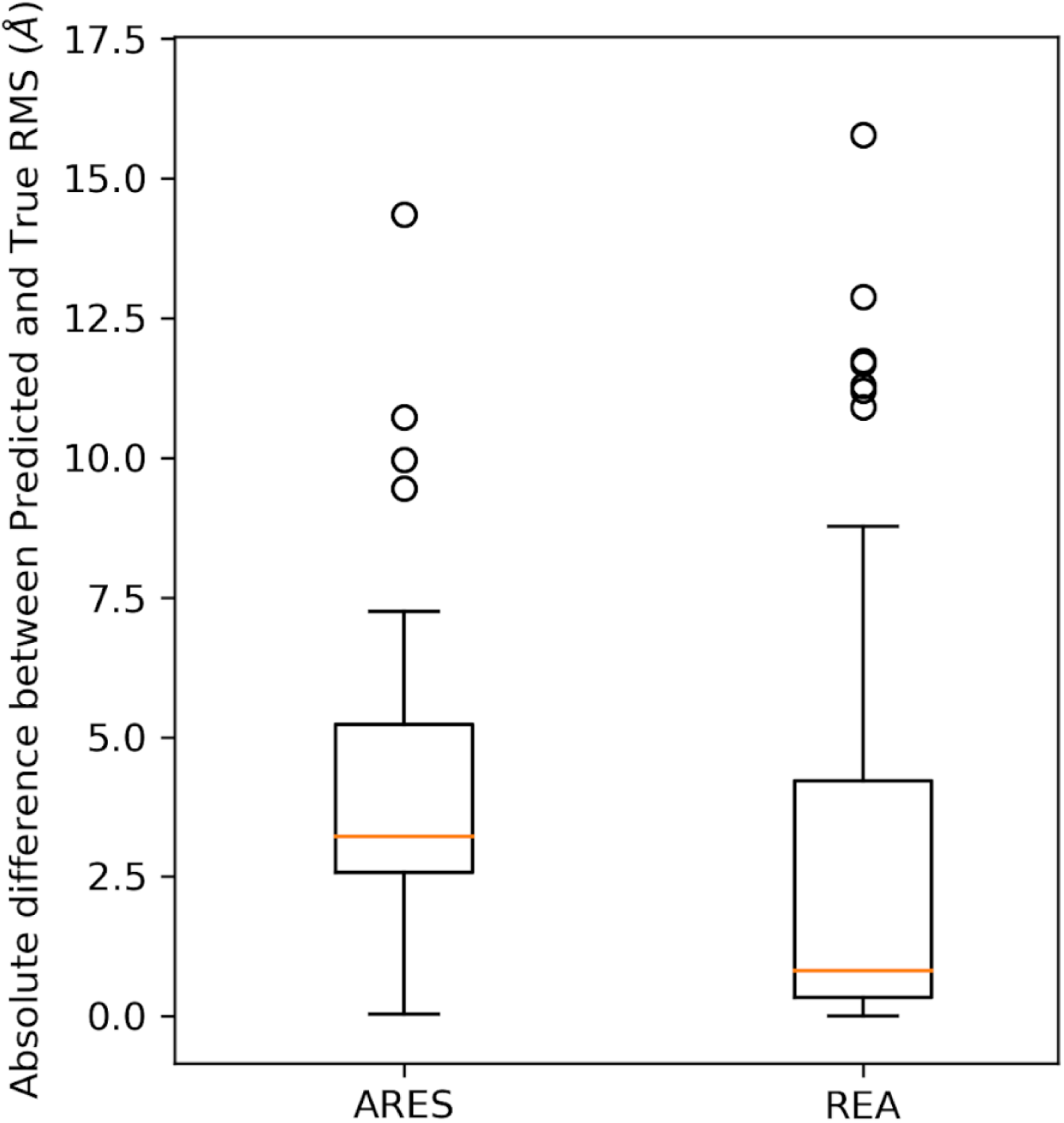
Boxplots of absolute differences between predicted and true RMSD for ARES and REA on pseudoknotted RNAs (58 RNA 3D structures).

Figure 14 shows that on the pseudoknotted test set the RMSD predictions made by REA have a lower difference from the true RMSD values than the RMSD predictions made by ARES. The two-sided Wilcox signed-rank test was conducted to compare the absolute differences between true RMSD and the RMSD predicted by REA and the absolute difference between true RMSD and the RMSD predicted by ARES. The test revealed a significant difference between the two sets of predictions (p = 0.0064). We might assume that this is due to the fact that REA was trained only on pseudoknotted structures. This will be addressed below.

We compare REA to ARES based on the correlation between the predicted and true RMSD. Figure 15 presents the regression plots of predicted RMSD against the true RMSD for ARES and REA on the pseudoknotted test set, along with the Pearson’s correlation coefficient for each plot.

**Figure 15:**
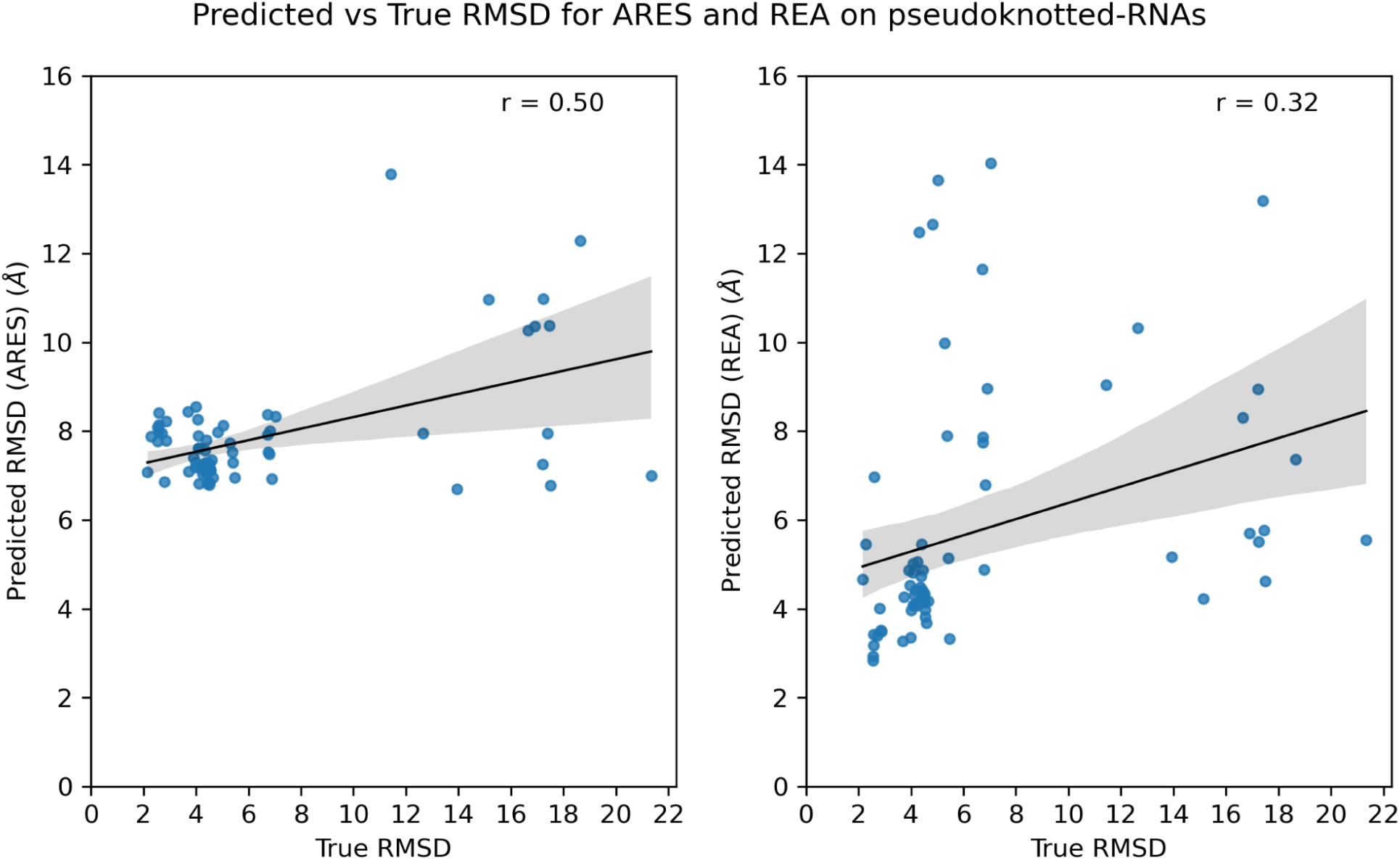
Comparison of predictions by ARES and REA on pseudoknotted RNAs. The figure displays scatter plots of the true RMSD values on the x-axis and the corresponding predictions made by ARES (left) and REA (right) on the y-axes. Each scatter plot is accompanied by a fitted line representing the linear regression model fit. Both models exhibit a positive linear relationship with the true RMSD values, with close values for Pearson’s correlation coefficient.

Based on the results shown in Figure 15, on the pseudoknotted test set, the RMSD predicted by ARES is more positively correlated to the true RMSD than the RMSD predicted by REA. On the test set with pseudoknotted RNAs, ARES predicts a lower RMSD for the less accurate structures and a higher RMSD for the most accurate structures. On the contrary, REA predicts some of the more accurate structures (True RMSD ∼ 5 Å) to be less accurate [Predicted RMSD > 8 Å]. Both REA and ARES underpredict the less accurate structures in the set (True RMSD > 15 Å).

### Model evaluation on non-pseudoknotted RNAs

After having evaluated REA on pseudoknotted RNAs, we evaluated REA on non-pseudoknotted RNAs as well (Supplementary, Table 3). Evaluating REA on non-pseudoknotted RNAs, while it was trained on a pseudoknotted-only dataset, gives insights on how models can generalize on different types of 3D RNA structures. We evaluate REA in a similar way as we did on the pseudoknotted RNAs. Figure 16 shows the distribution of the absolute differences between the predicted and the true RMSD for ARES, REA and the baseline on the non-pseudoknotted RNAs.

**Figure 16:**
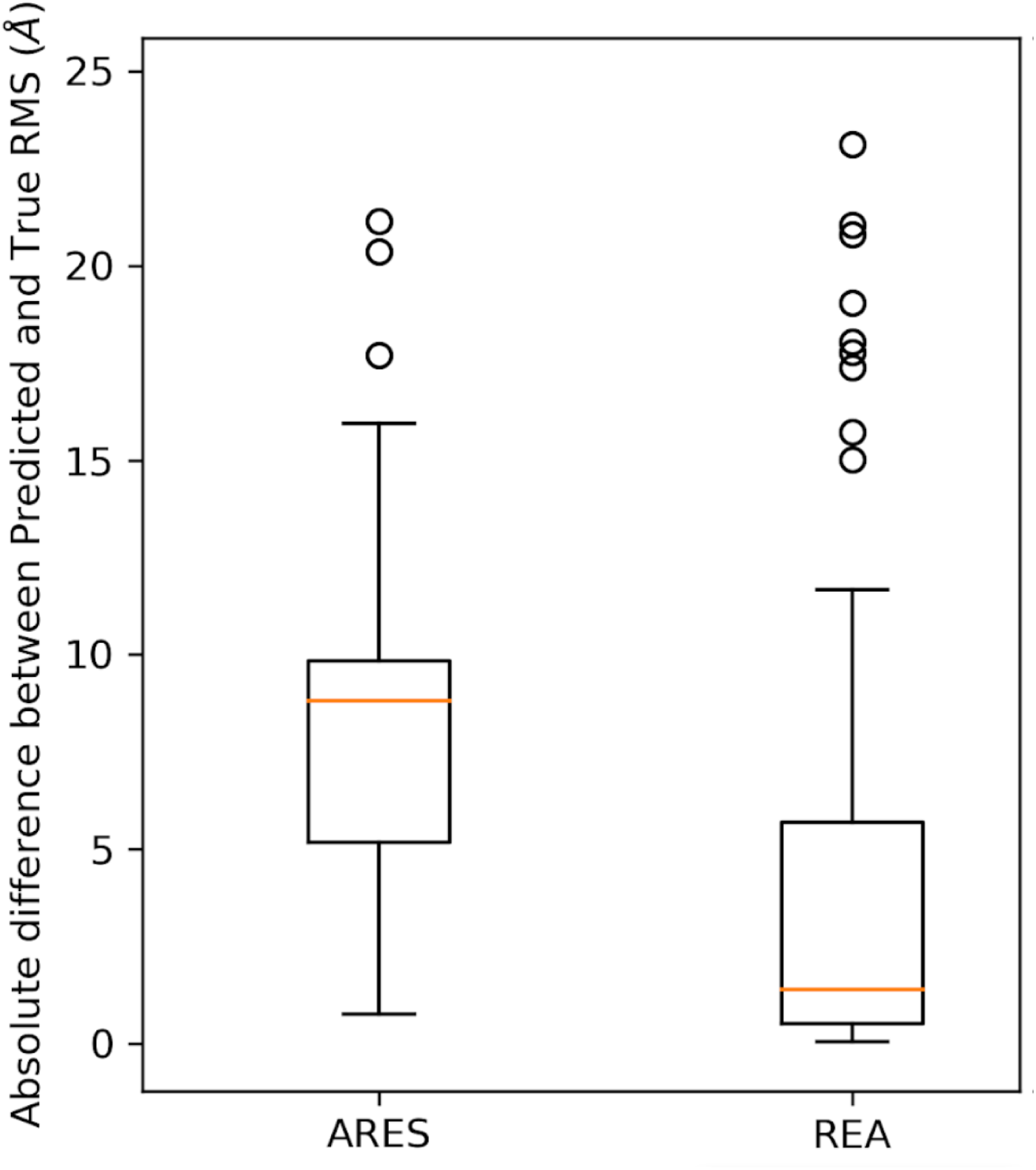
Boxplots of absolute differences between predicted and true RMSD for ARES andREA on non-pseudoknotted RNAs (117 RNA 3D structures). ARES shows the largest differences. REA exhibits the smallest median difference, indicating the most accurate predictions.

Figure 16 shows that on the non-pseudoknotted test set the RMSD predictions made by REA have a lower difference from the true RMSD values than the RMSD predictions made by ARES, as it was noticed on the pseudoknotted test set. The two-sided Wilcox signed-rank test was conducted to compare the absolute differences from true RMSD of the predictions made by REA and the absolute difference from true RMSD of the predictions made by ARES on the non-pseudoknotted test set. The test revealed a significant difference between the two sets of predictions at significance level 5% (W = 1162, p = 4.75e-10).

Additionally, we compare REA to ARES on the correlation between the predicted and true RMSD, as we did for the pseudoknotted test set. Figure 17 presents the regression plots of predicted RMSD against the true RMSD for ARES and REA on the non-pseudoknotted test set, along with the Pearson’s correlation coefficient for each plot.

**Figure 17:**
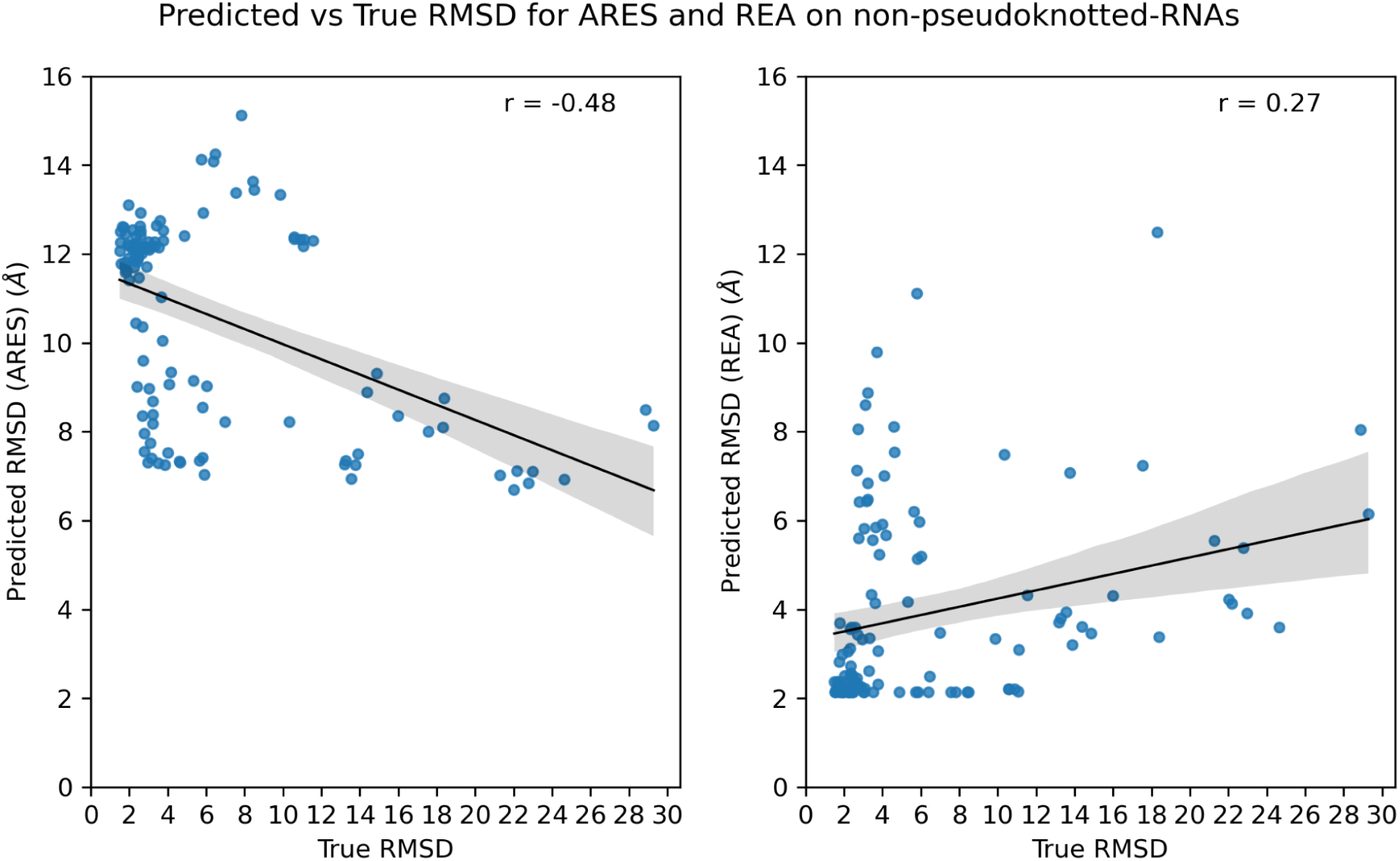
Comparison of predictions by ARES and REA on non-pseudoknotted RNAs. The figure displays scatter plots of the true RMSD values on the x-axis and the corresponding predictions made by ARES (left) and REA (right) on the y-axes. Each scatter plot is accompanied by a fitted line representing the linear regression model fit. ARES predictions exhibit a negative linear relationship with the true RMSD values, while REA predictions exhibit a slightly positive linear relationship with the true RMSD values.

Based on the results shown in Figure 17, on the non-pseudoknotted test set, the RMSD predicted by REA is positively correlated to the true RMSD, while the RMSD predicted by ARES is negatively correlated to the true RMSD. REA predicts more accurately the structures with small true RMSD than ARES, which incorrectly predicts many highly accurate structures to have an RMSD around 12Å. Starting from a high RMSD prediction for the accurate structures, ARES predicts a lower RMSD for the less accurate structures [Predicted RMSD (ARES) ∼ 8 Å]. REA outperforms ARES in the comparison of Pearson’s correlation coefficients, because it shows positive correlation and seems to have captured the variations in true RMSD more effectively.

### Optimal predictor for model selection

Lastly, we searched among the methods REA, ARES, SVR and the summation of their predictions (SumReaAres, SumReaSVR, SumAresSVR and SumReaAresSVR) for the method that would predict the most accurate model among the DeepFoldRNA 1, DeepFoldRNA 2, DeepFoldRNA 3, DeepFoldRNA 4, DeepFoldRNA 5, DeepFoldRNA 6, trRosettaRNA, RhoFold unrelaxed and RhoFold relaxed alternative models of an RNA chain. We predicted the RMSD of the models of a novel validation set with REA, ARES, SVR and their summations and selected the model with the lowest predicted RMSD (or the sum of the predicted RMSD) as the prediction of each selective prediction method. We compared the RMSD prediction methods based on the average true RMSD of the predicted models. We assessed the average true RMSD of the best prediction method against the best single model.

The validation set included 261 representatives from the Representative Set of RNA structures version 3.306 with a resolution of smaller or equal than 2.5 Å (NR set) [35]. The representatives in the NR Set are RNA chains that are non-redundant in structure, except homologous RNAs among species [35]. We cleaned the NR set by removing the common RNA chains between the nrlist representatives and RNA-Puzzles and then selecting the representatives with length greater or equal than 20 and less than 180 nucleotides.

As a test set we used the RNA-Puzzles set with the 39 RNAs (https://www.rnapuzzles.org/results/). The PDB files of the RNAs in RNA-Puzzles were downloaded from the standardized dataset (https://github.com/RNA-Puzzles/standardized_dataset) for the first 21 RNA-Puzzles, and from the Protein Data Bank (PDB) with the PDB codes for the last 29 RNA-Puzzles (Supplementary, Table 4). We splitted the PDBs of RNA-Puzzles into chains, extracted the RNA sequences from the PDB files and predicted the 3D structure of RNAs. For the RNA chains with common sequence (rp11A/rp11B, rp02B/rp02D, rp02F/rp02H, rp02A/rp02C/rp02E, rp33A/rp33B, rp36C/rp36D, rp37A/rp37B), we kept the best predicted RMSD among the alternative chains for each model (DeepFoldRNA 1, DeepFoldRNA 2, DeepFoldRNA 3, DeepFoldRNA 4, DeepFoldRNA 5, DeepFoldRNA 6, trRosettaRNA, RhoFold unrelaxed and RhoFold relaxed). The test set also included 4 chains with pseudoknots, 2WW9_F, 3IZF_C, 4UJN_J, 4UK0_B and 4UKN_J, that were excluded from the training and validation datasets. In total, there were 55 chains in our test set.

Table 1 shows the average true RMSD of the predicted models by the different RMSD prediction methods on the validation and test. REA has the best (minimum) average True RMSD in the validation set and the summation of REA, ARES and SVR predictions (SumReaAresSVR) has the best average true RMSD on the test set. Table 2 shows the average true RMSD of the models predicted by a single method on the validation and the test sets. The trRosettaRNA model has the best average true RMSD among single models on both validation and test set. Models predicted with a selective prediction method outperform trRosettaRNA.

**Table 1:** The average true RMSD of the models predicted by the different RMSD selective prediction methods on the validation set (NR Set) and the test set (RNA-Puzzles).

| RMSD Prediction Method | Average True RMSD of Predicted Model of NR Set | Average True RMSD of Predicted Model of RNA-Puzzles |
| --- | --- | --- |
| REA | **5.564** | 6.847 |
| ARES | 7.293 | 8.947 |
| SVR | 6.455 | 10.450 |
| SumReaAres | 6.068 | 6.784 |
| SumReaSVR | 5.721 | 6.784 |
| SumAresSVR | 6.324 | 8.744 |
| SumReaAresSVR | 5.704 | **6.690** |

**Table 2:**
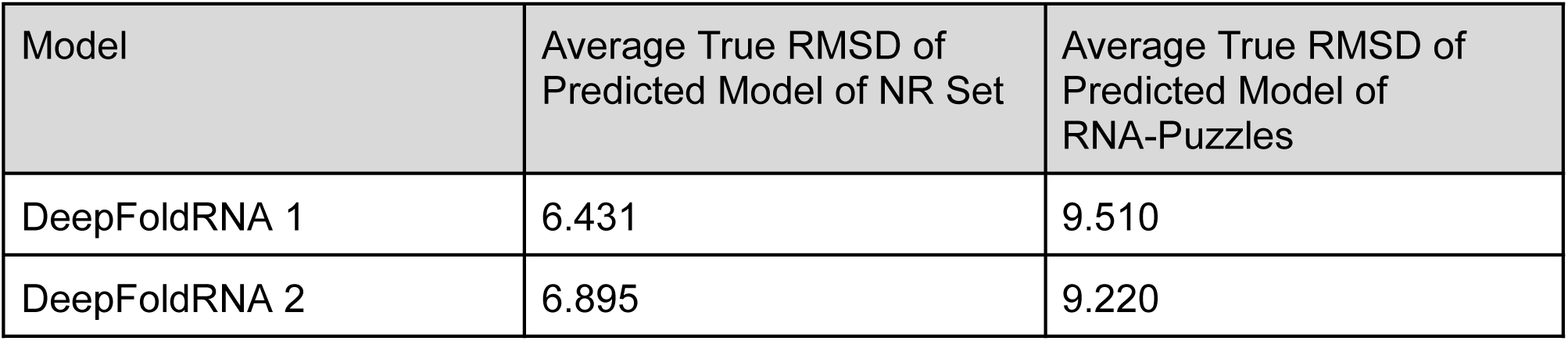
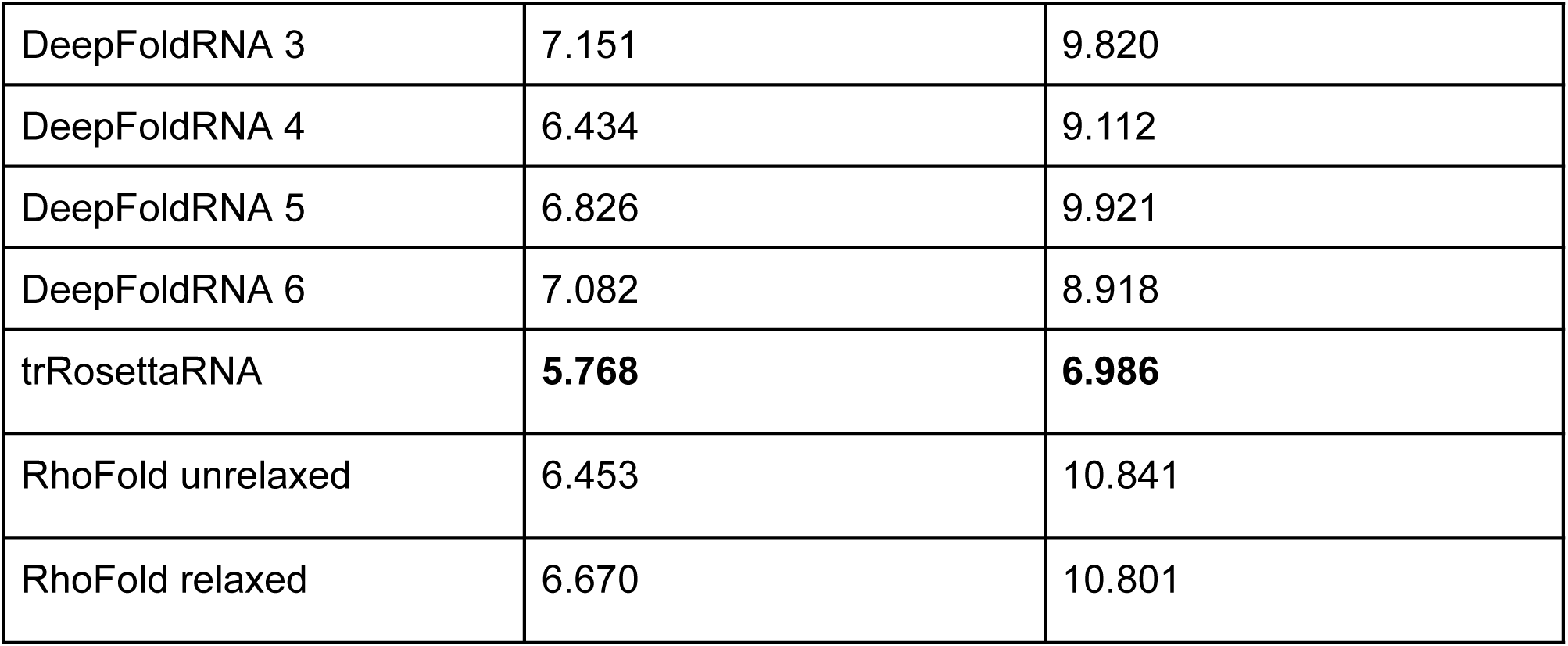
The average true RMSD of the models predicted by a single method on the validation set (NR Set) and the test set (RNA-Puzzles).

| Model | Average True RMSD of Predicted Model of NR Set | Average True RMSD of Predicted Model of RNA-Puzzles |
| --- | --- | --- |
| DeepFoldRNA 1 | 6.431 | 9.510 |
| DeepFoldRNA 2 | 6.895 | 9.220 |
| DeepFoldRNA 3 | 7.151 | 9.820 |
| DeepFoldRNA 4 | 6.434 | 9.112 |
| DeepFoldRNA 5 | 6.826 | 9.921 |
| DeepFoldRNA 6 | 7.082 | 8.918 |
| trRosettaRNA | **5.768** | **6.986** |
| RhoFold unrelaxed | 6.453 | 10.841 |
| RhoFold relaxed | 6.670 | 10.801 |

To assess the significance of this improved performance, bootstrapping was performed for the comparison of medians between the true RMSD of a method and the true RMSD of trRosettaRNA, for all RMSD prediction methods. Resampling was performed from the differences trRosettaRNA true RMSD - method true RMSD, 1. in our validation set and 2. in our test set, 1000000 times. A subsample of size 17 was used for both datasets. The p-value was calculated as the ratio of the times that the median of the sample is greater than the median of the original dataset: #(median_diff_sample > median_diff_dataset) / 1000000. Table 3 presents the p-values for each method on both validation and test set. The method with the best (minimum) p-value on both validation and test set is SumReaSVR, indicating an equal or better performance than trRosettaRNA alone.

Figure 18 shows the scatterplots of the True RMSD of the model predicted by SumReaSVR against the True RMSD of the trRosettaRNA model. The majority of the predictions made by SumReaSVR coincide with the trRosettaRNA model. In the test set, there are 3 cases for which SumReaSVR outperforms trRosettaRNA and none for the reverse.

**Figure 18:**
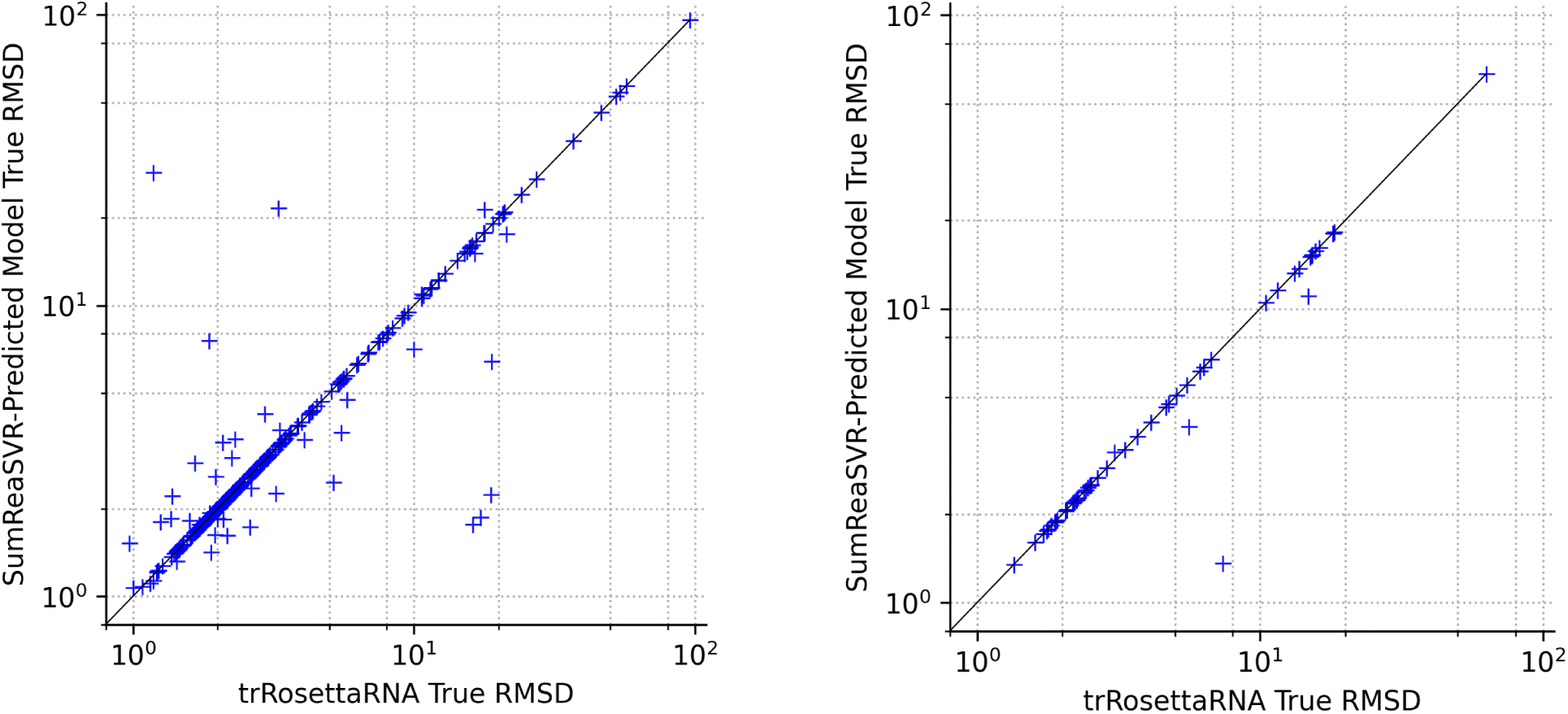
True RMSD of the SumReaSVR-predicted model vs. True RMSD of trRosettaRNA on the NR set (left) and RNA-Puzzles (right).

**Table 3:** The p-value, calculated with bootstrapping, for the difference of medians of True RMSD of each RMSD prediction method and the True RMSD of the trRosettaRNA model, on the validation set (NR Set) and the test set (RNA-Puzzles).

| RMSD Prediction Method | p-value NR Set | p-value RNA-Puzzles |
| --- | --- | --- |
| REA | 0.999999 | 0.000019 |
| ARES | 0.869653 | 0.801692 |
| SVR | 0.908467 | 0.967059 |
| SumReaAres | 0.991380 | 0.000544 |
| SumReaSVR | **0.000004** | **0.000000** |
| SumAresSVR | 0.954165 | 0.977884 |
| SumReaAresSVR | 0.999971 | 0.000093 |

Figure 19 shows an example where SumReaSVR predicted a more accurate structure for the RNA-Puzzle 2 than the trRosettaRNA model, with an RMSD of 6 Å smaller.

**Figure 19:**
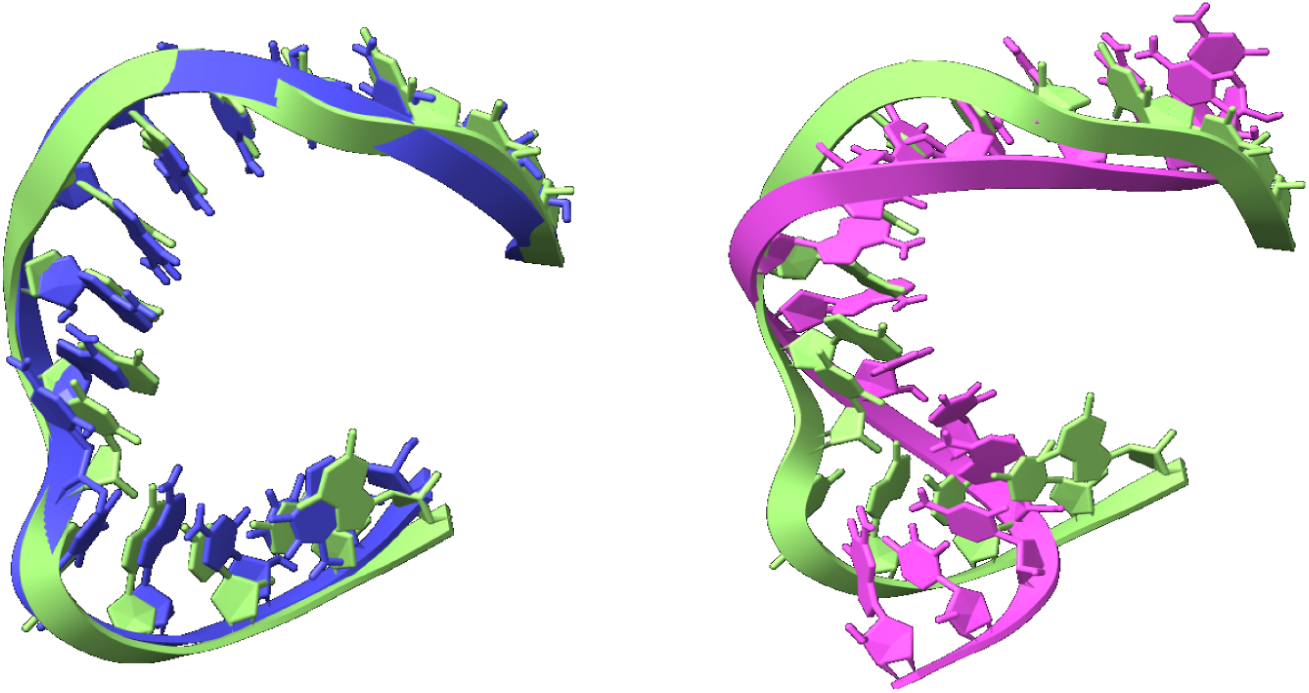
DeepFoldRNA 4 model (left, purple, RMSD = 1.37Å) and trRosettaRNA model (right, pink, RMSD = 7.38Å) for the RNA-Puzzles 2, chain A. Each model is superimposed to the experimentally determined structure (PDB: 3P59, chain A, green). The DeepFoldRNA 4 model was predicted by SumReaSVR.

## Conclusion

We have developed REA (RMSD Estimation Algorithm) which predicts the RMSD of a 3D RNA structure from its experimentally determined structure, using the stereochemical evaluation of the structure provided via Molprobity. REA was trained on a training pseudoknotted RNAs and tested on pseudoknotted and non-pseudoknotted test sets separately. Based on the absolute difference between the predicted and true RMSD, REA outperformed ARES on both pseudoknotted and non-pseudoknotted test sets. The combination of REA with ARES and an SVR model, trained and tuned on the same feature set as REA, led to seven RMSD prediction methods, from which the combination of REA and SVR predictions (SumReaSVR) significantly outperformed the best direct prediction method, trRosettaRNA.

Future work might focus on further optimizing the hyperparameters such as training for more than 300 epochs, testing values for the learning rate 0.02 and higher and values for the number of hidden units between 20 and 30. Also the use of some of the second order features shown to have high absolute correlation could be investigated. A comprehensive workflow that will process a given RNA sequence, call both the RNA structure prediction methods used in this work as well as promising novel methods such as epRNA [36] and rank all obtained structure models would be of high relevance for experimentalists.

## Data and Code Availability

The python code of the model and the training and test data are available at: https://github.com/ReaKal/RMSD-Estimation-Algorithm

## Acknowledgements

This work was funded by the Commissioned Service 2022-EIP1 “Alignment of the Interoperability Platform FAIR Service Architecture Framework with the Data Platform, Communities, and ELIXIR Projects” from ELIXIR, the research infrastructure for life-science data.

