## Supplemental material for "A Root Mean Square Deviation Estimation Algorithm (REA) and its use for improved RNA Structure Prediction"

This supplementary material contains 13 tables and 5 figures.

**Table 1: List of 26 RNA chains used for training REA.**

| |
|---|
| 2WWQ_V |
| 3AMU_B |
| 3J2L_3 |
| 3TS0_U |
| 3TS2_V |
| 3WQY_C |
| 4QK8_A |
| 4UJG_B |
| 4UJI_J |
| 4UJL_B |
| 4UJV_B |
| 4UJX_J |

| |
|---|
| 4UK2_J |
| 4UK5_B |
| 4UKC_J |
| 4UKG_B |
| 4UKI_J |
| 4UKQ_B |
| 4UKS_J |
| 4UKV_B |
| 4UKX_J |
| 4UL5_B |
| 4ULC_J |
| 4ULF_B |
| 4ULK_B |
| 4ULN_J |

**Table 2: List of RNA chains with pseudoknots used for testing REA.**

| |
|---|
| 2WW9_F |
| 3IZF_C |
| 4UJN_J |
| 4UK0_B |
| 4UKN_J |
| 4UL7_J |
| RNA-Puzzle 17 (chain A) * |
| RNA-Puzzle 18 (chain A) * |
| RNA-Puzzle 21 (chain A) * |

| |
|---|
| RNA-Puzzle 5 (chain A) * |
| RNA-Puzzle 8 (chain A) * |

* Downloaded from https://github.com/RNA-Puzzles/standardized_dataset

**Table 3: List of RNA chains without pseudoknots used for testing REA.**

| |
|---|
| RNA-Puzzle 2 (chain A) * |
| RNA-Puzzle 2 (chain B) * |
| RNA-Puzzle 2 (chain C) * |
| RNA-Puzzle 2 (chain D) * |
| RNA-Puzzle 2 (chain E) * |
| RNA-Puzzle 2 (chain F) * |
| RNA-Puzzle 2 (chain G) * |
| RNA-Puzzle 2 (chain H) * |
| RNA-Puzzle 4 (chain A) * |
| RNA-Puzzle 6 (chain A) * |
| RNA-Puzzle 7 (chain A) * |
| RNA-Puzzle 10 (chain B) * |
| RNA-Puzzle 13 (chain A) * |
| RNA-Puzzle 15 (chain A) * |
| RNA-Puzzle 19 (chain A) * |
| RNA-Puzzle 19 (chain B) * |
| RNA-Puzzle 20 (chain A) * |
| RNA-Puzzle 20 (chain B) * |

* Downloaded from https://github.com/RNA-Puzzles/standardized_dataset

**Table 4: RNA chains from RNA-Puzzles and pseudoknots dataset used for testing the different model selection methods.**

| PDB Code | Name in Text |
|---|---|
| 2WW9_F | 2WW9_F |
| 3IZF_C | 3IZF_C |
| 4UJN_J | 4UJN_J |
| 4UK0_B | 4UK0_B |
| 4UKN_J | 4UKN_J |
| 3MEI_A | rp01A |
| RNA-Puzzle 1 (chain B) * | rp01B |
| RNA-Puzzle 2 (chain A) * | rp02A |
| RNA-Puzzle 2 (chain B) * | rp02B |
| RNA-Puzzle 2 (chain C) * | rp02C |
| RNA-Puzzle 2 (chain D) * | rp02D |
| RNA-Puzzle 2 (chain E) * | rp02E |
| RNA-Puzzle 2 (chain F) * | rp02F |
| RNA-Puzzle 2 (chain G) * | rp02G |
| RNA-Puzzle 2 (chain H) * | rp02H |
| 3OWZ_A | rp03A |
| RNA-Puzzle 4 (chain A) * | rp04A |
| RNA-Puzzle 5 (chain A) * | rp05A |
| RNA-Puzzle 6 (chain A) * | rp06A |
| RNA-Puzzle 7 (chain A) * | rp07A |
| RNA-Puzzle 8 (chain A) * | rp08A |

| | |
|---|---|
| 5KPY_A | rp09A |
| RNA-Puzzle 10 (chain A) * | rp10A |
| RNA-Puzzle 10 (chain B) * | rp10B |
| RNA-Puzzle 11 (chain A) * | rp11A |
| RNA-Puzzle 11 (chain B) * | rp11B |
| RNA-Puzzle 12 (chain A) * | rp12A |
| RNA-Puzzle 13 (chain A) * | rp13A |
| RNA-Puzzle 14 bound (chain A) * | rp14Abound |
| RNA-Puzzle 14 free (chain A) * | rp14Afree |
| RNA-Puzzle 15 (chain A) * | rp15A |
| RNA-Puzzle 15 (chain B) * | rp15A |
| 6Y0Y_A | rp16A |
| RNA-Puzzle 17 (chain A) * | rp17A |
| RNA-Puzzle 18 (chain A) * | rp18A |
| RNA-Puzzle 19 (chain A) * | rp19A |
| RNA-Puzzle 19 (chain B) * | rp19B |
| RNA-Puzzle 20 (chain A) * | rp20A |
| RNA-Puzzle 20 (chain B) * | rp20B |
| RNA-Puzzle 21 (chain A) * | rp21A |
| 6JQ5_A | rp22A |
| 6JQ5_B | rp22B |
| 6E8U_B | rp23B |
| 6OL3_C | rp24C |

| | |
|---|---|
| 6P2H_A | rp25A |
| 6PMO_B | rp26B |
| 6POM_A | rp27A |
| 6POM_B | rp27B |
| 6UFM_A | rp28A |
| 6UFM_B | rp28B |
| 6TB7_A | rp29A |
| 7BG9_B | rp30B |
| 7MLX_R | rp31R |
| 7EOJ_A | rp32A |
| 7ELP_A | rp33A |
| 7ELP_B | rp33B |
| 7VE9_A | rp34A |
| 7QR4_B | rp35B |
| 7QR3_C | rp36C |
| 7QR3_D | rp36D |
| 8GXC_A | rp37A |
| 8GXC_B | rp37B |
| 8HB8_A | rp38A |
| 8DP3_R | rp39R |

* Downloaded from https://github.com/RNA-Puzzles/standardized_dataset

**Table 5: Median MSE CV and IQR MSE CV of combinations of Feature Sets, Learning Rate and Maximum Epochs grouped by Learning Rate and Feature Set. Ordered by ascending Median MSE CV. The combination chosen as the optimal is highlighted in bold.**

| Feature set | Learning Rate | Median MSE CV | IQR MSE CV |
|---|---|---|---|
| **Molprobity** | **0.001** | **4.3907** | **4.9676** |
| Molprobity | 0.0002 | 5.4558 | 5.7164 |
| Molprobity | 0.0001 | 7.3361 | 11.2481 |
| Molprobity | 5E-05 | 9.7145 | 39.8052 |
| Molprobity+Energies | 0.001 | 53.1605 | 255.3255 |
| Molprobity+Energies | 0.0002 | 136.0152 | 20576.0401 |
| Molprobity+Energies | 0.0001 | 16007.4336 | 448615.6562 |
| Molprobity+Energies | 5E-05 | 138722.4297 | 1925907.2129 |

**Table 6: Median MSE CV and IQR MSE CV of combinations of Feature Sets, Learning Rate and Maximum Epochs grouped by Number of Hidden Units and Feature Set.**

| Feature Set | Number of Hidden Units | Median MSE CV | IQR MSE CV |
|---|---|---|---|
| **Molprobity** | **15** | **5.0755** | **5.3679** |
| Molprobity | 10 | 5.9855 | 6.1288 |
| Molprobity | 5 | 8.2905 | 22.6502 |
| Molprobity+Energies | 15 | 147.0672 | 16624.5685 |
| Molprobity+Energies | 10 | 494.9626 | 137380.9838 |
| Molprobity+Energies | 5 | 5782.5256 | 16624.5685 |

**Table 7: Median MSE CV and IQR MSE CV of combinations of Feature Sets, Learning Rate and Maximum Epochs grouped by Number of Maximum Epochs and Feature Set.**

| Feature Set | Number of maximum epochs | Median MSE CV | IQR MSE CV |
|---|---|---|---|
| **Molprobity** | **300** | **5.5780** | **5.9818** |
| Molprobity | 150 | 7.2872 | 13.0398 |
| Molprobity+Energies | 300 | 161.3546 | 39037.4966 |
| Molprobity+Energies | 150 | 3130.2340 | 390718.7466 |

**Table 8: Median MSE CV and IQR MSE CV of different learning rates (first learning rate refinement).**

| Learning Rate | Median MSE CV | IQR MSE CV |
|---|---|---|
| 0.0128 | 2.4197 | 6.1610 |
| 0.0096 | 2.4997 | 4.2324 |
| **0.0012** | **2.5045** | **3.3541** |
| 0.0048 | 2.5581 | 5.6142 |
| 0.0024 | 2.5862 | 5.3731 |
| 0.0006 | 2.5880 | 2.8840 |
| 0.0016 | 2.6954 | 4.6508 |
| 0.0008 | 2.7209 | 3.0890 |
| 0.0032 | 2.7551 | 3.5295 |
| 0.0064 | 2.9104 | 4.6850 |
| 0.0004 | 3.2798 | 6.7420 |
| 0.0002 | 4.3332 | 9.4805 |
| 0.0001 | 4.5663 | 14.0062 |
| 5E-05 | 11.0691 | 25.8936 |

**Table 9: Median MSE CV and IQR MSE CV of different numbers of hidden units.**

| Number of Hidden Units | Median MSE CV | IQR MSE CV |
|---|---|---|
| 9 | 2.5028 | 4.4885 |
| 17 | 2.5298 | 3.1356 |
| 11 | 2.5300 | 3.2676 |
| 19 | 2.5488 | 4.5083 |
| **12** | **2.6034** | **3.0088** |
| 16 | 2.6901 | 3.7164 |
| 10 | 2.7481 | 5.0226 |
| 20 | 2.7696 | 3.9572 |
| 13 | 2.7823 | 3.8221 |
| 14 | 2.8077 | 3.7159 |
| 18 | 2.8652 | 3.2486 |

| | | |
|---|---|---|
| 15 | 2.9080 | 3.7605 |
| 8 | 3.3804 | 5.9172 |
| 7 | 3.3905 | 6.8614 |
| 6 | 3.9577 | 7.9214 |
| 5 | 4.9184 | 6.2498 |

**Table 10: Median MSE CV and IQR MSE CV of different learning rates (second learning rate refinement).**

| Learning Rate | Median MSE CV | IQR MSE CV |
|---|---|---|
| 0.0096 | 1.2500 | 3.7184 |
| **0.0064** | **1.5224** | **3.0782** |
| 0.0048 | 1.8278 | 3.7430 |
| 0.0006 | 1.9393 | 5.2713 |
| 0.0024 | 1.9476 | 4.0314 |
| 0.0012 | 2.0598 | 3.7848 |
| 0.0128 | 2.0734 | 3.7243 |
| 0.0032 | 2.0762 | 4.2906 |
| 0.0016 | 2.3020 | 3.7267 |
| 0.0008 | 2.3168 | 4.0448 |
| 0.0002 | 2.6524 | 5.0419 |
| 0.0004 | 2.7292 | 4.0823 |
| 0.0001 | 5.4792 | 11.3947 |
| 5E-05 | 8.0578 | 22.7151 |

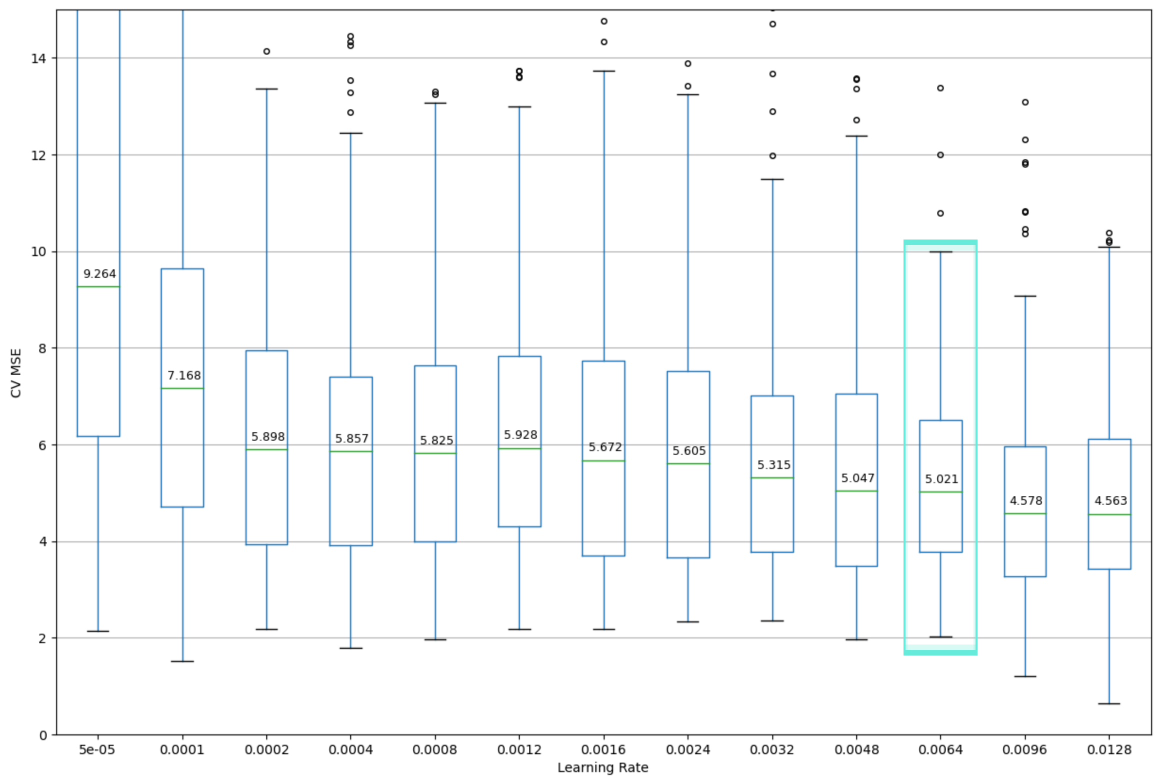


**Figure 1: Cross-validation performance of different learning rates with 20 iterations and the initial Molprobity feature set. The rest of the hyperparameters remain fixed (n_units = 12, max_epochs = 300, batch_size = 32).**

**Table 11: Median MSE CV and IQR MSE CV of different learning rates (second learning rate refinement, initial Molprobity feature set, 20 iterations).**

| Learning Rate | Median MSE CV | IQR MSE CV |
|---|---|---|
| 0.0128 | 4.5625 | 2.6979 |
| 0.0096 | 4.5775 | 2.6788 |
| **0.0064** | **5.0207** | **2.7324** |
| 0.0048 | 5.0466 | 3.5615 |
| 0.0032 | 5.3149 | 3.2432 |
| 0.0024 | 5.6052 | 3.8554 |
| 0.0016 | 5.6725 | 4.0355 |
| 0.0008 | 5.8245 | 3.6415 |
| 0.0004 | 5.8569 | 3.4907 |
| 0.0002 | 5.8980 | 4.0195 |
| 0.0012 | 5.9281 | 3.5171 |
| 0.0001 | 7.1684 | 4.9234 |
| 5E-05 | 9.2635 | 11.1015 |

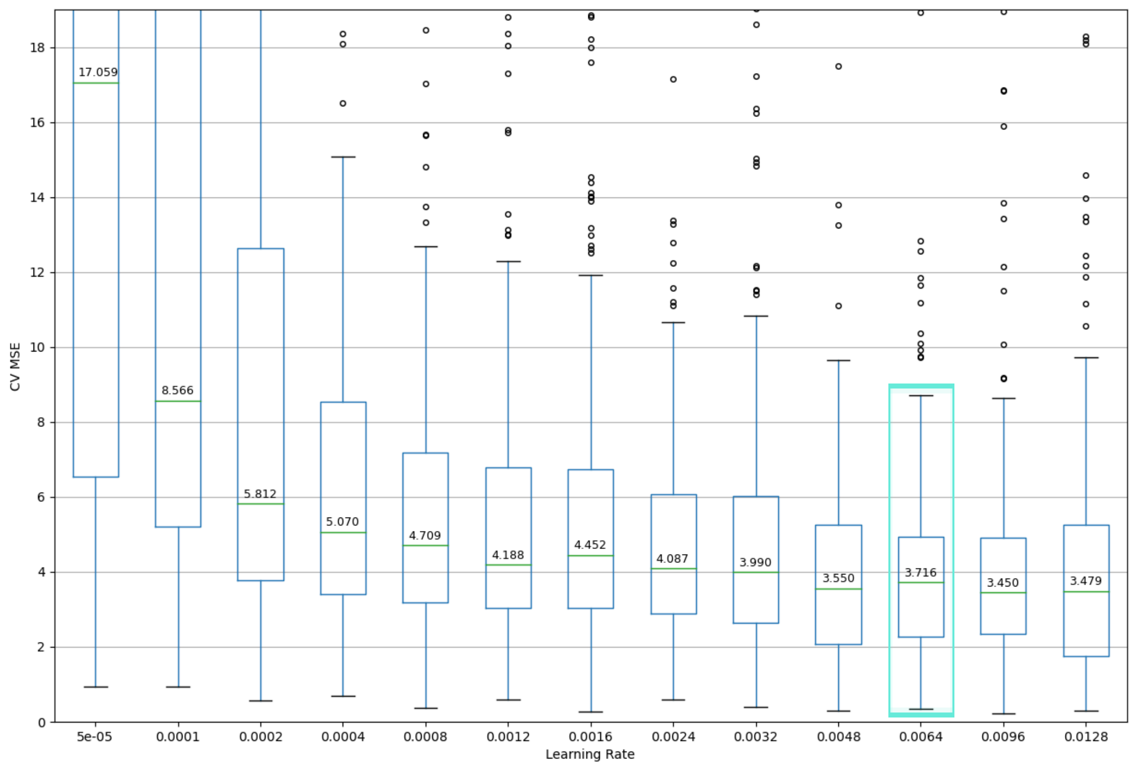


**Figure 2: Cross-validation performance of different learning rates with 20 iterations and the enhanced Molprobity feature set. The rest of the hyperparameters remain fixed (n_units = 12, max_epochs = 300, batch_size = 32).**

**Table 12: Median MSE CV and IQR MSE CV of different learning rates (second learning rate refinement, enhanced Molprobity feature set, 20 iterations).**

| Learning Rate | Median MSE CV | IQR MSE CV |
|---|---|---|
| 0.0096 | 3.4500 | 2.5717 |
| 0.0128 | 3.4793 | 3.4874 |
| 0.0048 | 3.5504 | 3.1822 |
| **0.0064** | **3.7163** | **2.6595** |
| 0.0032 | 3.9898 | 3.3643 |
| 0.0024 | 4.0871 | 3.1977 |
| 0.0012 | 4.1881 | 3.7590 |
| 0.0016 | 4.4520 | 3.7016 |
| 0.0008 | 4.7089 | 4.0035 |
| 0.0004 | 5.0700 | 5.1470 |
| 0.0002 | 5.8117 | 8.8523 |
| 0.0001 | 8.5658 | 14.8853 |
| 5E-05 | 17.0585 | 74.6341 |

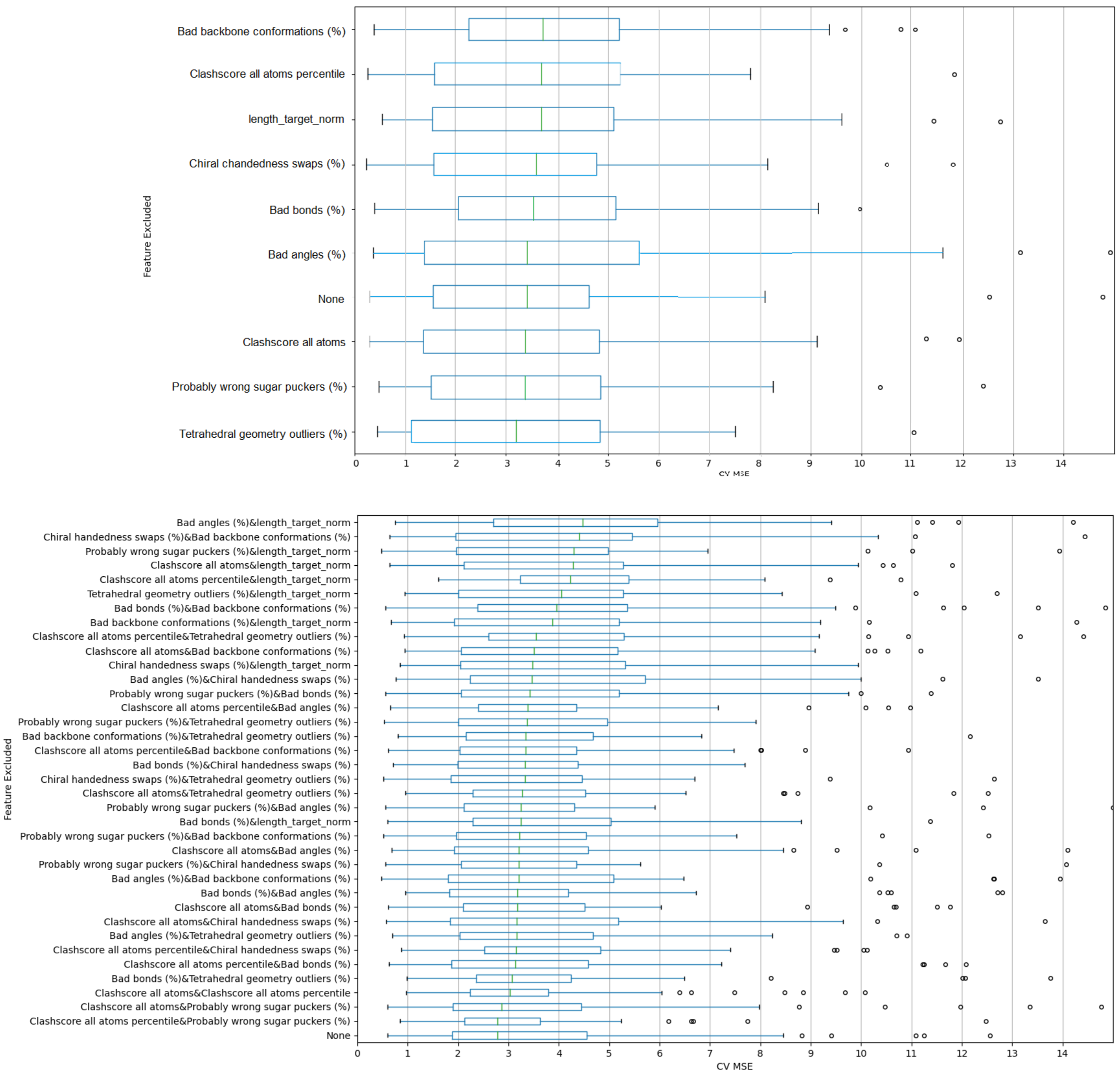


**Figure 3: Cross-validation (10-fold with 5 iterations) performance of features subsets where different features were knocked out.**

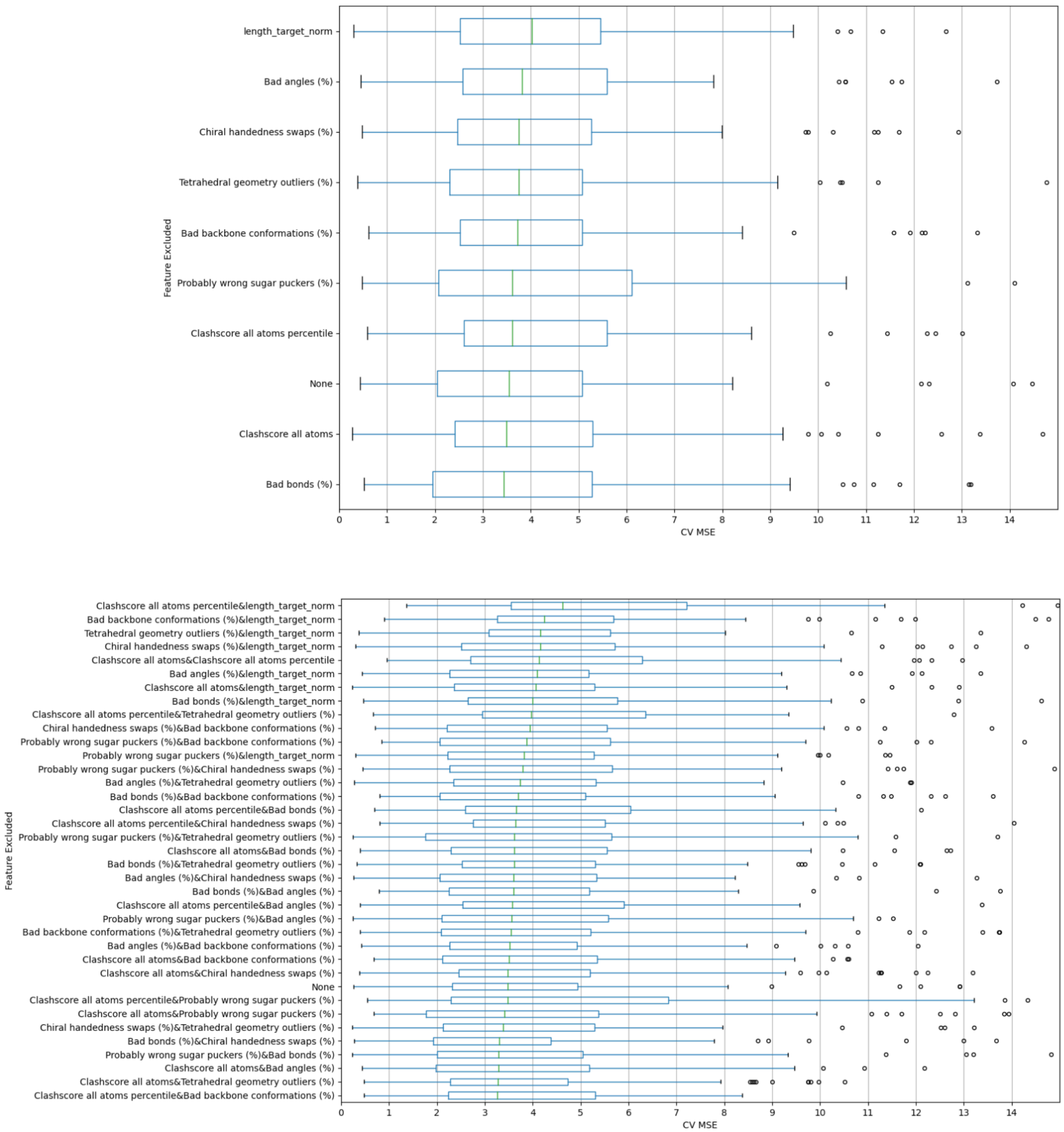


**Figure 4: Cross-validation (10-fold with 10 iterations) performance of features subsets where different features were knocked out.**

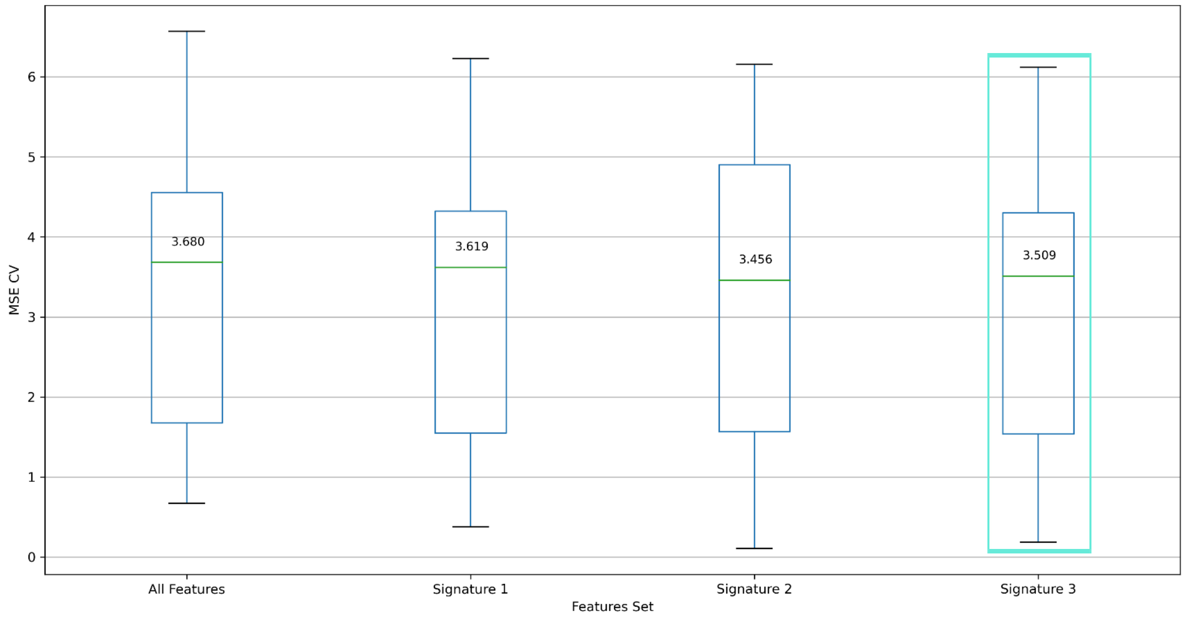


**Figure 5: Cross-validation performance of the SVR model with the total feature set and the three different signature sets found by JADBio.**

**Table 13: Median MSE CV and IQR MSE CV for different feature sets for the SVR model.**

| Feature set | Median MSE CV | IQR MSE CV |
|---|---|---|
| All features | 3.680 | 2.879 |
| Signature 1 | 3.619 | 2.771 |
| Signature 2 | 3.456 | 3.335 |
| Signature 3 | **3.509** | **2.768** |
